# Farnesyltransferase inhibition overcomes the adaptive resistance to osimertinib in *EGFR*-mutant NSCLC

**DOI:** 10.1101/2022.04.01.486707

**Authors:** Sarah Figarol, Célia Delahaye, Rémi Gence, Raghda Asslan, Sandra Pagano, Claudine Tardy, Jacques Colinge, Jean-Philippe Villemin, Antonio Maraver, Irene Ferrer, Luis Paz-Ares, Isabelle Lajoie-Mazenc, Estelle Clermont, Anne Casanova, Anne Pradines, Julien Mazières, Olivier Calvayrac, Gilles Favre

**Author notes:** Equal contribution (Sarah Figarol, Célia Delahaye, Olivier Calvayrac, Gilles Favre).

## Abstract

Drug-tolerant “dormant” cells (DTC) have emerged as one of the major non-genetic mechanisms driving resistance to targeted therapy in lung cancer, although the sequence of events leading to entry and exit from dormancy remain poorly described. Here, we performed real-time monitoring of the cell cycle dynamics during the adaptive response to Epidermal Growth Factor Receptor tyrosine kinase inhibitors (EGFR-TKi) in a panel of EGFR-mutated lung cancer cell lines. We identified a rare population of S/G2 cycling cells (referred to as early escapers) that emerged in the first hours of treatment amongst stably arrested and progressively dying G1 cells. We determined that early escapers evolved from a non-proliferative differentiated alveolar type 1 (AT1) phenotype which was invariably associated with cytoskeletal remodeling through Rho/ROCK pathway activation. Using a panel of Rho-pathway inhibitors, we found that the farnesyltransferase inhibitor tipifarnib induced a complete clearance of EGFR-TKi-induced DTC thus fully preventing relapse *in vitro*. Using a xenograft model and a PDX model of EGFR^L858R/T790M^ lung cancer, co-treatment with tipifarnib prevented relapse to osimertinib for up to 6 months with no evidence of toxicity. Among the farnesylated proteins regulated during osimertinib treatment, concomitant inhibition of RHOE and LaminB1 was sufficient to recapitulated FTIs’ effect. Osimertinib and tipifarnib co-treatment completely suppressed the emergence of AT1 phenotype, prevented mitosis of S/G2-treated cells and increased the apoptotic response through activation of ATF4-CHOP-dependent Integrated Stress Response (ISR) pathway. Our data strongly support the use of tipifarnib in combination with osimertinib in patients to effectively and durably prevent relapse.

## RESULTS

### Drug-tolerance is a dynamic rather than a dormant state

Although an increasing number of studies have focused on characterization of drug-tolerant “dormant” cells (DTC) and/or fully resistant proliferative clones (RPC)^1-6^, the kinetics of evolution throughout the different states is largely unknown. We used the FUCCI (fluorescence ubiquitination cell cycle indicator) system^7^ to perform a real time monitoring of cell cycle dynamics in response to 1µM EGFR-TKi erlotinib or osimertinib in a panel of *EGFR*-mutated cell lines. Importantly, cell lines were previously subcloned to minimize/avoid the presence of potential pre-existing resistant cells^2,8^ (Supp Fig. 1A and B). For all cell lines, we observed a common pattern of G1 accumulation within the first 48 hours of treatment (Fig. 1a, Supp Fig. 2A, videos), which was invariably associated with p27^Kip1^ overexpression and Retinoblastoma (Rb) protein dephosphorylation (Supp Fig. 2B). While most cells remained in G1 and progressively died resulting in bulk population decrease, a subset of cells (referred to as early escapers) rapidly progressed though S/G2 (Fig. 1a-c, Supp. Fig. 2A, videos), an observation consistent with a rare, drug-tolerant cycling subpopulation as recently described^9^. Interestingly, although DTC and early escapers could be observed in H3255- and HCC2935-treated cells, they failed to form resistant proliferative clones in these conditions (Supp Fig 3A-D) but retained the capacity to resume proliferation after drug withdraw (Supp Fig. 3E).

**Figure 1:**
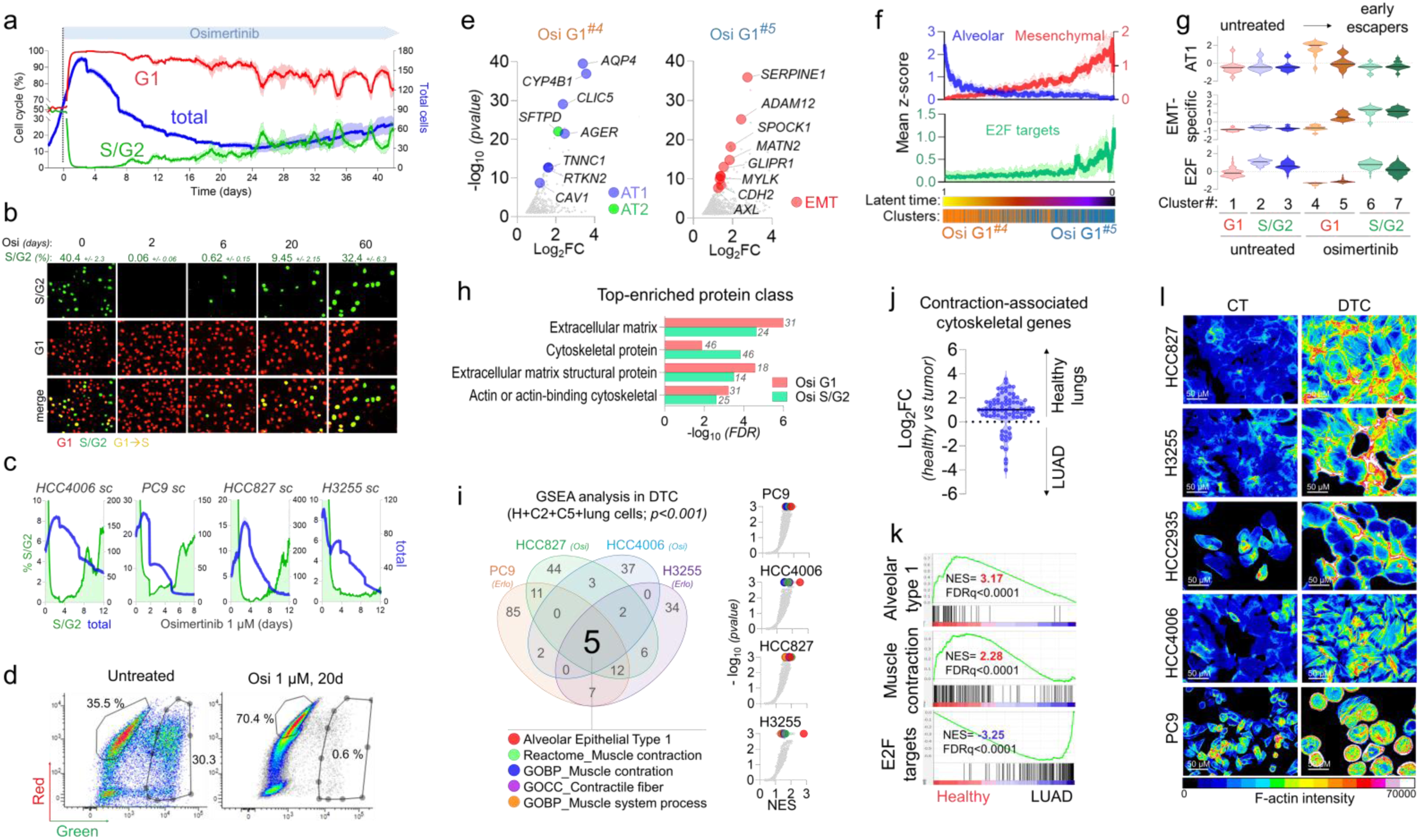
Drug-tolerance is a dynamic state which invariably implicates alveolar differentiation and cytoskeletal remodeling. a: Percentage of total cells (blue), S/G2 (green) or G1 (red) population of HCC4006 subclonal cells during osimertinib treatment (1μM). b: Representative fluorescence images of S/G2 (green) or G1 (red) HCC4006 subclonal cells during osimertinib treatment (1 μM). c: S/G2 (green) or G1 (red) fractions (%) of PC9, HCC4006, HCC827, HCC2935, H3255 subclonal cells during osimertinib treatment (1 μM). d: Flow cytometry charts of red cells (G1) or green cells (S/G2 or G1 -S) of untreated or osimertinib treated (1μM, 20 days) HCC4006 subclonal cells sorted for single-cell RNA sequencing experiment. Percentages of each fractions according to total population are specified. e: Osimertinib-induced genes overexpressed specifically in G1 cluster 4 vs cluster 5 (left) or in cluster 5 vs cluster 4 (right). AT 1 (blue), AT2 (green) or EMT (red)-specific genes are highlighted. f: Mean z-score evolution of alveolar (blue), mesenchymal (red) and E2F_target (green) signatures according to latent time cell repartition. Osi-G1 cluster 4 (orange) and cluster 5 (blue) are specified. g: Violin plots representing the mean z-score values of alveolar type 1 (AT 1), EMT-specific and E2F_target signatures in the different clusters. h: Top significantly overrepresented protein classes (pantherbd.org) in osimertinib-treated HCC4006 G1 (red bars) and S/G2 cells (green bars) compared to untreated-G1 and S/G2 populations, respectively (log2FC>1, p-value<0.05) i: Left: Venn diagram comparing the significantly enriched pathways (p<0.001) in erlotinib- or osimertinib-derived drug-tolerant cells. GSEA analysis was performed on differentially expressed genes (p<0.05) obtained by RNAseq in PC9 and H3255 cells were treated with 1 μM erlotinib for 21 days, HCC827 cells treated with 1 μM osimertinib for 11 days, and HCC4006 G1 -cells treated with osimertinib for 21 days, compared to respective control cells. Right: Volcano plots of the pathways enriched in indicated EGFR-TKi-treated DTC. j: Differential expression of the contraction-associated cytoskeletal genes in healthy lungs compared to lung adenocarcinoma using TCGA data. k: GSEA analysis of alveolar type 1, muscle-contraction and E2F_targets signatures in healthy lungs compared to lung adenocarcinoma using TCGA data. l: Phalloidin F-actin staining of HCC827, H3255, HCC2935, HCC4006 and PC9 control cells or DTC upon osimertinib treatment (1μM) (X20 magnification, scale bar = 50μM). Images were analyzed by Image J software (filter: 16 colors).

These results suggest that the establishment of an adaptive resistance mechanism through a drug-tolerant state is a dynamic and progressive process and is highly variable even among genetically homogeneous cellular models.

### Early escapers emerge from a differentiated alveolar-like phenotype with contractile features

To characterize the molecular mechanisms underlying evolution from G1-arrest to early escape, we performed scRNAseq on ∼3000 G1 (red) and S/G2 (green) cells sorted from both untreated and osimertinib-treated HCC4006 cells, which enabled enrichment of the rare population of early escapers (Fig. 1d, Supp Fig. 4A and B). In line with previous reports^2,4^, Myogenesis and Epithelial-to-Mesenchymal (EMT) signatures were strongly upregulated in both G1- and S/G2-treated cells, while cell cycle-related gene signatures such as E2F_targets or G2M_checkpoint were profoundly downregulated in G1 and fully restored in early escapers (Supp Fig. 4C). We observed that the most deeply cell cycle-repressed cluster (Osi G1^*cluster #4*^, Supp Fig. 4B) was highly enriched in alveolar type 1 (AT1)-related genes such as *AQP4, CYP4B1, CLIC5, AGER* or *TNNC1*, but also in alveolar type 2 (AT2)-specific marker *SFTPD* (Surfactant Protein D) (Fig. 1e, Supp Fig. 4D), consistent with recent data reported in minimal residual disease (MRD) of osimertinib-treated NSCLC patients^10^. Interestingly, these genes were among the most overexpressed in healthy lungs compared with adenocarcinomas as revealed by a differential expression analysis using the TCGA database (Supp Fig. 4E). Pseudotime analysis revealed a progressive switch from an alveolar to a mesenchymal transcriptomic profile in the Osi G1^*cluster #5*^ (Fig. 1e and f, Supp Fig. 5A-C) which was further enhanced in early escapers (Fig. 1g, Supp Fig. 5D) strongly suggesting that the EMT process could have been initiated from the alveolar-like sub-population. Using recently published gene signatures of the different lung cell types^11^, we observed that lineage switching involved a robust downregulation of mucous/goblet-related genes such as *MUC5B, BPIFB2, SCGB3A1* or *AZGP1* as well as a strong overexpression of fibroblast and muscle-specific genes (Supp Fig. 5E, Supp Fig. 6A-C). This process was strongly associated with extracellular matrix- and muscle-specific gene signatures (Supp Fig. 6D) and correlated with a strong enrichment in cytoskeletal- and extracellular matrix-coding genes (Fig. 1h), with a different pattern of expression during the phenotypic switch (Supp Fig. 6E). Since these features could be specific to the HCC4006 model, we performed RNAseq experiments in drug-tolerant cells generated from other *EGFR*-mutated cell lines treated with either erlotinib or osimertinib at 1µM. Although each cell line seemed to follow a different path of differentiation (Supp Fig. 7A-C), all cell lines consistently overexpressed alveolar type 1 and contractile-related gene signatures (Supp Fig. 7D and E), which were the unique common gene signatures among the models tested (Fig. 1i). Interestingly, gene signatures related to cellular contractility were highly specific to the drug-tolerant state and did not necessarily evolve toward a mesenchymal phenotype (Supp Fig. 8). Indeed, H3255-derived DTC strongly overexpressed alveolar and muscle-specific genes but were not associated with EMT signatures (Supp Fig. 9A) and did not express EMT-associated genes such as *SERPINE1, AXL, SPOCK1*, or EMT-TF (Supp Fig. 9B), which could explain their inability to form fully resistant proliferative clones. Strikingly, a similar enrichment in contraction-associated cytoskeletal genes and muscle specific signatures was observed in healthy lungs compared to lung adenocarcinomas (Fig. 1j and k, Supp Fig. 10A-D), strongly reinforcing the evidence that the DTC subtype shared phenotypic resemblance with normal epithelial lung cells. Using our transcriptomic data, we built a new signature of drug tolerance composed of 350 genes that were commonly upregulated in our models (*i*.*e*. log2FC>1 in at least 4 out of 5 cell lines) (Supp Fig. 11A). This signature was positively associated with osimertinib-induced drug-tolerance in a model of osimertinib-treated EGFR-mutated lung PDX (see Fig. 4) and in residual disease in patients (Supp Fig. 11B) but also with healthy lungs compared to lung adenocarcinomas (LUAD) (Supp Fig. 11C and D). Finally, we aimed to assess the physiological consequences of this increased contractile signatures. We observed a highly reorganized cytoskeleton in all the DTC as revealed by F-actin staining, with the presence of actin stress fibers and/or lamelipodia, depending on the cell type (Fig. 1l, Supp Fig. 12 and 13). Interestingly, stress fibers were not detected in the H3255 cell line, which was in line with the absence of EMT signatures, however increased cortical F-actin was observed.

Taken together, our data suggest that osimertinib-resistant cells emerge from a differentiated drug-tolerant phenotype that shares similarities with an alveolar type 1 phenotype and involves cytoskeletal remodeling and contractile features.

### Farnesyltransferase inhibitors prevent relapse to EGFR-TKi *in vitro*

Given that the presence of F-actin often correlates with activation of one or more Rho-GTPases^12,13^, we hypothesize that Rho-proteins could be involved in the establishment of the drug-tolerant contractile state. Indeed, RNAseq data revealed that expression of *RHOB* and *RND3* (*RHOE*) was consistently increased in EGFR-TKi-derived DTC *in vitro* as well as in osimertinib-treated PDX (Fig. 2a). Using available *in vivo* scRNAseq data from Pasi Janne’s lab^4^, we observed that *RHOB* and *RND3* were among the most highly upregulated genes in PC9 xenografts treated with osimertinib or with osimertinib + trametinib (Fig. 2b). RHOB and RHOE were also increased at the protein level and correlated with a transient increase in phospho-MLC2 and phospho-cofilin, suggesting activation of the Rho/ROCK pathway during an early stage of drug-tolerance (Fig. 2c, Supp Fig. 14A). Interestingly, RHOB overexpression appeared to be highly associated with cell cycle arrest (Fig. 1c, Supp Fig. 14A-D) and its expression was also correlated with an AT1 signature in lung cancer patients as revealed by a TCGA analysis (Supp Fig. 14C). By contrast, RHOE expression was correlated with EMT and fibroblastic features (Supp Fig. 14B and E), suggesting different functions for the two GTPases during the adaptive response. These data are in agreement with our findings describing a role for RHOB in resistance to targeted therapy in EGFR-mutated lung cancer patients^14^ and BRAF-mutated melanoma^15^, although implication of RHOB or RHOE in the establishment of a drug-tolerant state has never been assessed. These observations prompted us to assess the efficacy of a panel of inhibitors of the Rho/ROCK pathway to prevent relapse to EGFR-TKi, such as ROCK inhibitors (Y27632, GSK269962A, RKI1447), the RhoA/B/C inhibitor C3-exoenzyme (tatC3), AKT inhibitor (G594), farnesyltransferase inhibitor (FTI, tipifarnib) and geranylgeranyltransferase inhibitor (GGTi, GGTi-298). Strikingly, all the inhibitors, excepted RKI1447, displayed cytotoxic or cytostatic activity on DTC and strongly delayed the emergence of resistant clones (Fig. 2d and e), however only tipifarnib fully prevented the development of resistances to osimertinib or erlotinib in all the cell line tested (Fig. 2d and e, Supp Fig. 15A), with no significant effect when used alone (Supp Fig. 15B). Combination with other FTIs such as CP-609754, lonafarnib or FTI-2153 showed similar effect, leading to cell death (Fig. 2f, Supp Fig. 15C). Interestingly, tipifarnib could also prevent the emergence of resistance to EGFR-TKi in both subclonal and parental cell lines (Fig. 2d-f), suggesting that inhibition of farnesyltransferase could also affect potentially pre-existing resistant subclones (Supp. Fig. 1A). Unexpectedly, treatment of early-emerging osimertinib-resistant proliferative clones with tipifarnib also lead to tumor regression, suggesting that these cells still displayed vulnerability to inhibition of farnesyltransferase (Fig. 2g and h). However, complete clearance of tumor cells was not often achieved when tipifarnib was used after relapse, suggesting that the cells that had acquired resistance to osimertinib displayed reduced sensitivity to farnesyltransferase inhibition (Fig. 2h, Supp Fig. 16).

**Figure 2:**
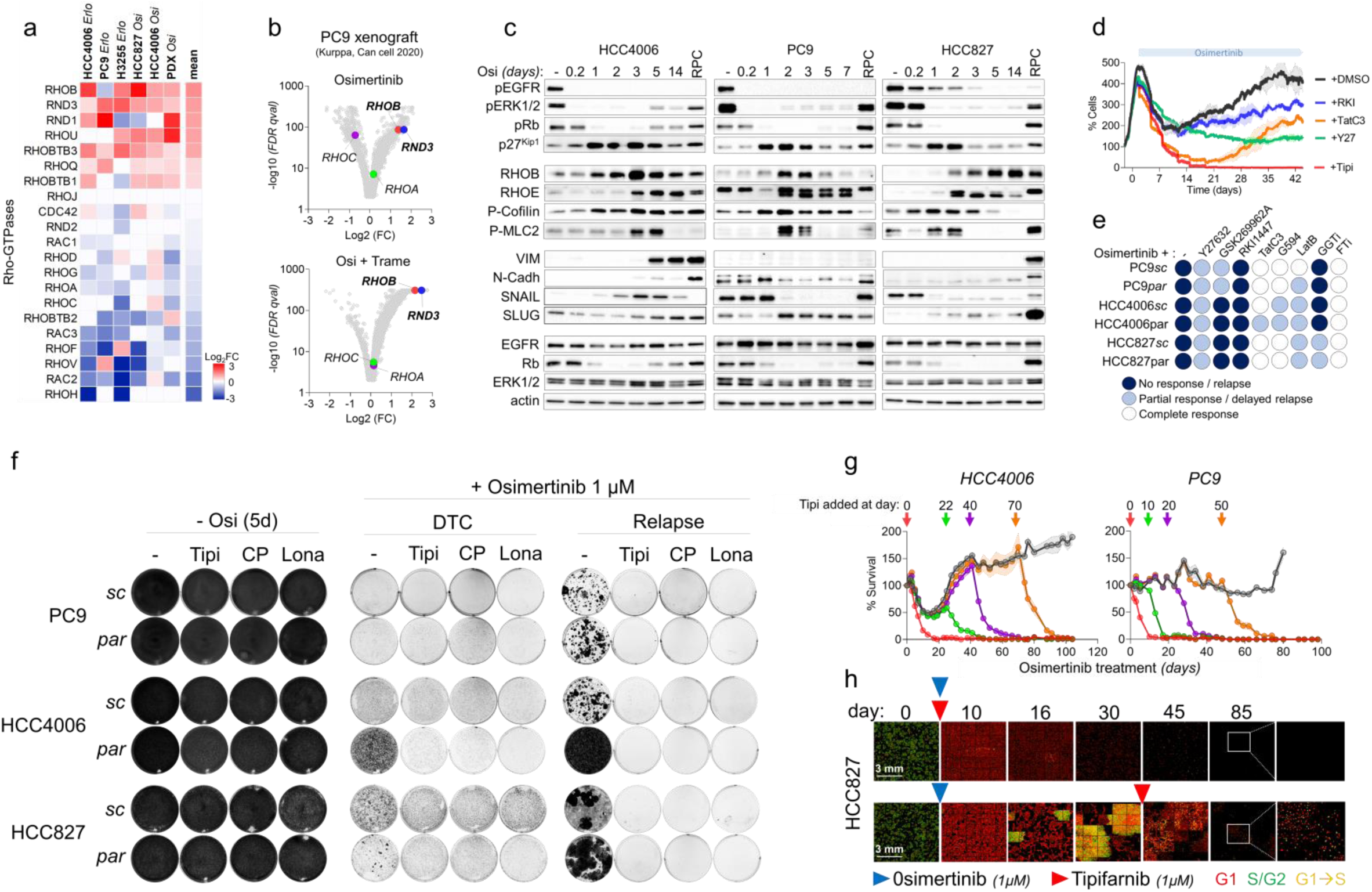
Farnesyltransferase inhibitors prevent resistance to EGFR-TKi *in vitro*. a: Differential mRNA expression of the 21 known Rho-GTPases in DTC generated from erlotinib-treated (HCC4006, PC9, H3255) or osimertinib-treated (HCC827, HCC4006) cells or in a PDX model of EGFR-mutated lung adenocarcinoma (described in Fig. 4). b: Volcano plot of the differentially expressed genes in osimertinib- (up) or osimertinib+trametinib- (down) treated PC9 xenografts. Data were obtained from a scRNAseq published by Kurppa et al. Cancer Cell 2020). c: Representative Western Blot of proteins related to EGFR pathway (phospho-EGFR, EGFR, phospho-ERK, ERK), cell contractility (phospho-MLC2, phospho-cofilin), EMT (vimentin, N-cadherin, SNAIL, SLUG), cell cycle (p27Kip1, phospho-RB, RB) and RhoGTPases (RHOB, RHOE) during osimertinib treatment (1μM) in HCC4006, PC9 and HCC827 subclonal cells. RPC= resistant proliferative clone. d: Cell survival (%) of HCC4006 parental cells during osimertinib treatment (1μM) alone or in combination with ROCK inhibitors (RKI, RKI1474 and Y27, Y27632), RHOA/B/C inhibitor (TatC3) or farnesyl-transferase inhibitor (Tipi, tipifarnib). e: Cell survival of PC9, HCC4006, HCC827 parental and subclonal cells upon osimertinib treatment (1μM) alone or in combination with ROCK inhibitors (Y27632, GSK269962A, RKI1447), RHOA/B/C inhibitor (TatC3), AKT inhibitor (G594), actin inhibitor (LatB, Latruncunlin B), geranylgeranyltransferase inhibitor (GGTI) or farnesyltransferase inhibitor (Tipi, tipifarnib). Deep blue round refers to no response or relapse, light blue round to partial or delayed response and white round to complete response. f: Crystal violet staining of clonal and parental PC9, HCC4006 and HCC827 cells treated for 5 days, at DTC state or after relapse with osimertinib (1μM) alone or in combination with farnesyltransferase inhibitors (tipifarnib 1μM, Tipi; CP-609.754 1μM, CP; lonafarnib 10μM, Lona). g: Cell survival (%) of HCC4006 and PC9 cells during osimertinib treatment (1μM) with or without tipifarnib (1μM) added at indicated timepoints. h: Images of HCC827 clonal cells upon osimertinib (1 μM) +/- tipifarnib (1μM). G1 cells are red-fluorescent, S/G2 cells are green-fluorescent and G1-S cells are in yellow. Blue and red arrows represent osimertinib treatment and tipifarnib treatment beginning respectively.

Our data show that Rho/ROCK pathway inhibitors efficiently prevent relapse to EGFR-TKi, among which tipifarnib displayed the most efficient and profound anti-tumor effect on both DTC and early-emerging clones.

### Osimertinib+Tipifarnib treatment induces mitotic defects and integrated stress response (ISR)-mediated apoptotic pathway

To decipher the molecular mechanisms underlying tipifarnib’s ability to prevent relapse to osimertinib, we performed scRNAseq on osimertinib + tipifarnib (OT)-treated G1 and S/G2 sorted cells and compared transcriptomes with those from cells treated with osimertinib. Strikingly, osimertinib-induced genes associated with our DRUG_TOLERANT_UP signature were drastically downregulated by tipifarnib as were many of the gene signatures related to EMT, myogenesis and cell-cycle (Fig. 3a and b, Supp Fig. 17A-D). Unexpectedly, tipifarnib prevented the upregulation of alveolar type 1 and fibroblast-specific genes but maintained muscle-associated genes such as *MYL7, ACTC1* or *ACTA1*, specifically in the S/G2 population (Fig. 3c, Supp Fig. 18A-C). We aimed to decipher the physiological consequence of tipifarnib treatment on DTC. Time-lapse imaging of FUCCI-transduced cells revealed that, although tipifarnib did not prevent the emergence of osimertinib-derived early escapers, escaping cells failed to undergo mitosis and ultimately died (Fig. 3d), suggesting that tipifarnib may interfere with cell division in these cells. We also observed that a high percentage of OT-treated cells that progressed to S/G2 returned to G1 without dividing, consistent with a process of mitotic slippage or endoreplication^16,17^, which was also observable at lower frequency in osimertinib-treated cells (Fig. 3d, Supp Fig. 19). Although this process is often used by cells as a mechanism of drug-resistance^16,17^, none of the cells that underwent this process were able to divide and ultimately died (Fig. 3d, videos). We also observed that OT-treatment strongly upregulated ATF4-regulated genes^18^ such as *DDIT3/CHOP, ATF3, CHAC1, PSAT1, FAM129A/NIBAN1* and the pro-apoptotic *BBC3/PUMA* (Fig. 3e and f). ATF4 can be induced by several processes including the Unfolded Protein Response (UPR) or the Integrated Stress Response (ISR)^19^. Upregulation of ATF4 and CHOP by OT treatment was confirmed at the protein level and correlated with PARP and Caspase 3 cleavage (Fig. 3g), strongly suggesting that activation of this pathway was responsible for OT-induced cell death.

**Figure 3:**
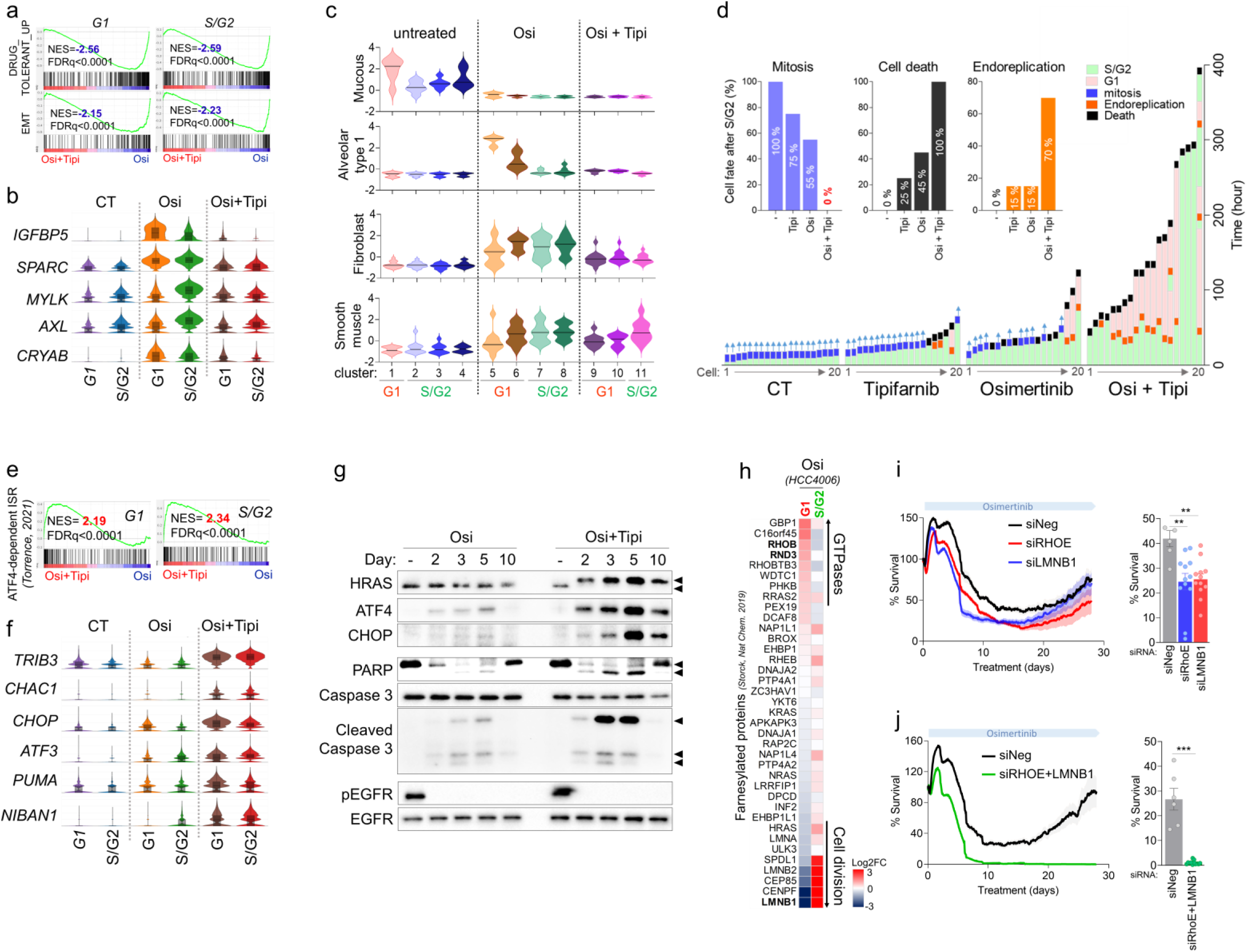
Tipifarnib causes multiple interferences in the adaptive response to osimertinib. a: GSEA analysis of Drug-Tolerant_Up and EMT signatures comparing Osimertinib+Tipifanib-versus Osimertinib-treated HCC4006 G1 and S/G2 cells. b: Violin plots representing the Log2 mRNA expression of drug-tolerant and EMT-related genes significantly altered by osimertinib+tipifarnib vs osimertinib treatment. c: Mean z-score of normal cell type-associated signatures (from Maynard, Nature 202010) significantly regulated by osimertinib and/or osimertinib+tipifarnib treatments. d: Cell fate after entry to S/G2 of HCC4006 clonal cells treated or not with osimertinib (1μM), tipifarnib (1 μM) or the combo. S/G2 cells are in green, G1 cells are in red. Mitosis event are represented in blue, endoreplication in orange and death in black. e: GSEA analysis of ATF4-dependent Integrated Stress Response (ISR) signature (Torrence, elife 202118) significantly modulated by Osimertinib+Tipifarnib versus Osimertinib treatment f: Violin plots representing the Log2 mRNA expression of ATF4-regulated genes significantly upregulated by Osimertinib+tipifarnib treatment g: Protein expression by western blot of proteins related to EGFR pathway (phospho-EGFR, EGFR), farnesyltransferase inhibitor efficacy (HRAS), apoptosis (PARP and caspase-3) and ISR (ATF4, CHOP) on HCC4006 clonal cells treated with osimertinib (1μM) alone or in combination with tipifarnib (1μM). h: Differential expression of genes coding for farnesylated proteins (Stock et al, Nat Chem, 2019) in osimertinib-treated G1 versus G1-untreated HCC4006 cells or osimertinib-treated S/G2 versus osimertinib-treated G1 cells i: Cell survival (%) of HCC4006 clonal cells during osimertinib treatment (1μM) and transfected with siRNA control (Neg) or targeting RHOE or LMNB1. j: Cell survival (%) of HCC4006 clonal cells during osimertinib treatment (1 μM) and transfected with siRNA control (Neg) or targeting RHOE and LMNB1.

### Identification of potential pharmacological targets of tipifarnib

We sought to identify farnesylated proteins that might explain tipifarnib’s ability to prevent relapse to EGFR-TKI. Consistent with previous findings^14^, siRNA-mediated inhibition of RHOB, but also RHOE, increased initial sensitivity to osimertinib (Supp Fig. 20A and B), although it failed to fully prevent relapse (Supp Fig. 20B, *data not shown*), suggesting that inhibition of these GTPases could not fully recapitulate tipifarnib’s effect. Using a recently published list of farnesylated proteins^20^ we observed in our scRNAseq data that several GTPases including *RHOB* and *RND3* were overexpressed by osimertinib in G1-treated cells, whereas the expression of cell-division-related genes such as *LMNB1, CENPF* or *SPDL1* was highly repressed during G1 but fully restored in S/G2 (Fig. 3h). Although individual inhibition of these genes increased sensitivity to osimertinib (Fig. 3i, Supp Fig. 20A and B), only inhibition of both RHOE and LaminB1 induced a complete clearance of osimertinib-treated cells, thus recapitulating tipifarnib’s effect (Fig. 3j), suggesting inhibition of both proteins was needed to fully prevent relapse to osimertinib. Collectively, our data suggest that tipifarnib prevents relapse to EGFR-TKi through a multi-targeted effect involving GTPases and cell-cycle regulators, resulting in impaired phenotypic differentiation, mitotic defaults and subsequent cell death through an ATF4-CHOP-dependent apoptotic pathway.

### Tipifarnib prevents relapse to EGFR-TKI *in vivo*

Finally, we aimed to test the combination of osimertinib and tipifarnib (OT) *in vivo* using several models of EGFR-mutated lung tumors. In a PC9 xenograft model, combination of osimertinib (5 mg/Kg, q.d., 5d/w) and tipifarnib (80 mg/Kg, b.i.d., 5d/w) induced a strong and stable response for up to 6 months with an almost complete response in all animals, whereas osimertinib induced a less potent effect and relapse was observed in 4/10 tumors (Fig. 4a-c, Supp Fig. 21A). OT treatment was well tolerated by the mice as indicated by a stable body weight, with no discernible effect on the general appearance of the mice (Supp Fig. 21B). We used the same treatment protocol in a PDX model of EGFR^L858R/T790M^ lung adenocarcinoma (TP103^21^). Osimertinib induced a moderate response with relapse of all (10/10) mice within 2 months, whereas OT-treatment induced significantly higher tumor regression in the first months with an almost stable disease lasting up to 5 months (Fig. 4d-g, Supp Fig. 22A-H), at which time OT-treated mice were sacrificed for subsequent molecular analysis of their tumors. As before, OT treatment was well tolerated by the mice as indicated by stable body weight (Supp Fig. 21B and F) and a good general aspect, thus confirming a good long-term tolerability of the treatment. Consistently with *in vitro* data, OT-treated tumors showed a strongly reduced proliferative state even after 5 months of treatment as highlighted by Ki67 staining (Fig. 4h). This correlated with increased p27^Kip1^ levels and reduced phospho-Rb as well as increased PARP and Caspase 3 cleavage, suggesting that non-cycling OT-treated cells were undergoing apoptosis (Fig. 4i). Low proliferation was confirmed by scRNAseq analysis (Fig. 4j-l, Supp Fig. 23), which also revealed an increased expression of ATF4-regulated genes (Fig 4m), which could suggest that the ISR pathway may also contribute to apoptosis *in vivo*.

**Figure 4:**
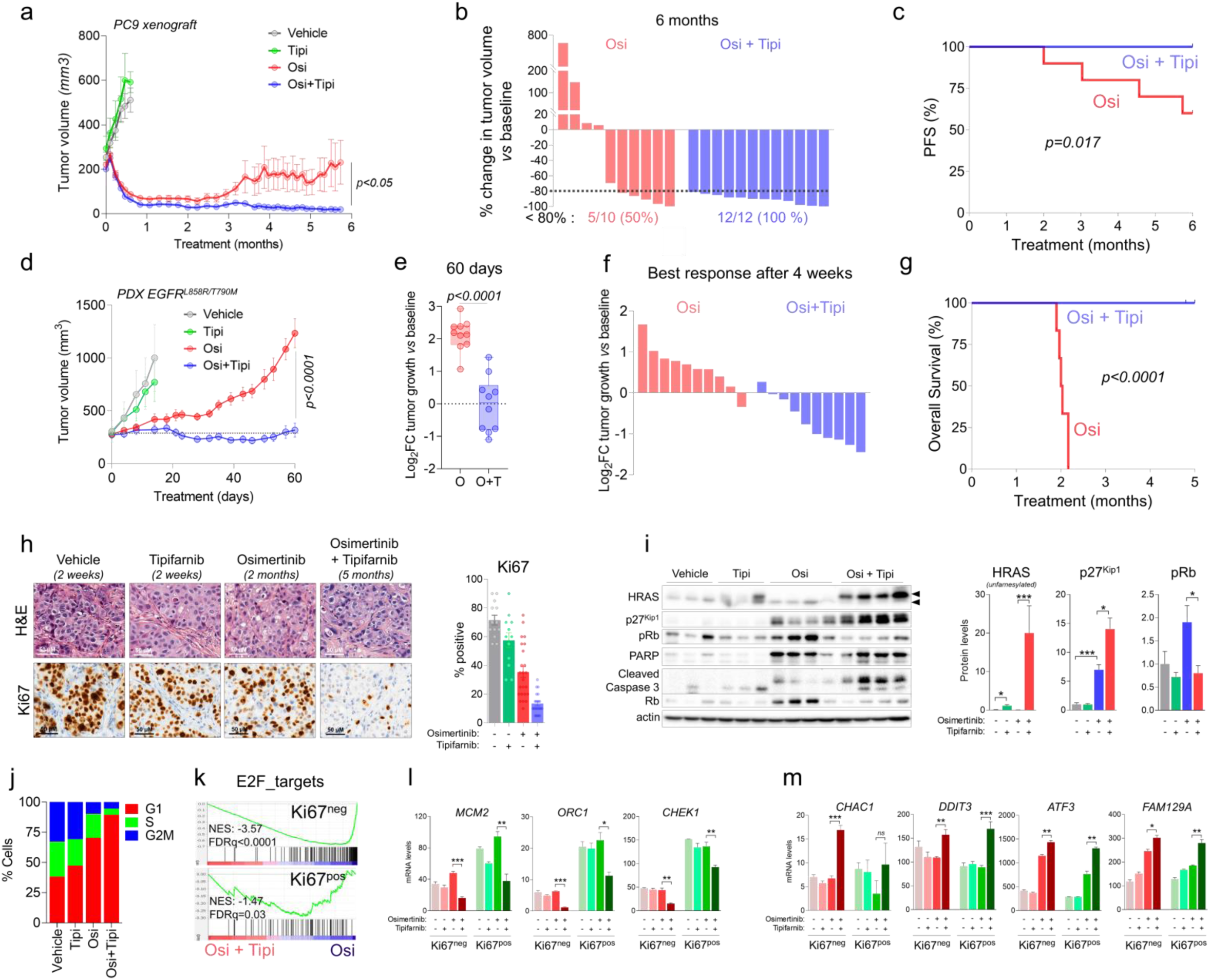
Tipifarnib prevents relapse to osimertinib *in vivo*. a: Mean tumor volume (mm3) of PC9 xenografts treated 5 days/week with vehicle, Tipifarnib (Tipi, 80mg/kg, b.i.d.), Osimertinib (Osi, 5 mg/kg, q.d), or by the combo (Osi + Tipi). n = 6 tumors in the vehicle and Tipifarnib arms, and n = 10 in the Osimertinib arm and n=12 in the combination arms. Mean ± SEM are shown. b: Change in tumor volume versus baseline of PC9 xenografts after 6 months of treatment with Osimertinib or a combination of Osimertinib and Tipifarnib. c: Progression-free survival of mice with PC9 xenografts and treated with Osimertinib or a combination of Osimertinib and Tipifarnib. P-value was determined by log-rank Mantel-Cox test. d: Mean tumor volume of an PDX model of EGFRL858R/T790M lung adenocarcinoma treated 5 days/week with vehicle, Tipifarnib (Tipi, 80mg/kg, b.i.d.), Osimertinib (Osi, 5 mg/kg, q.d), or by the combo (Osi + Tipi). The graph is the result of two independent cohorts of mice with a total of n = 4 tumors in the vehicle arm, n = 5 tumors in the Tipifarnib arm and n = 10 tumors in the Osimertinib and combination arms. Mean ± SEM are shown. e: Log2 fold change of the PDX growth compared to baseline after 60 days of treatment with Osimertinib or a combination of Osimertinib and Tipifarnib. f: Log2 fold change of the PDX growth compared to baseline at the best response from 4 weeks after treatment with Osimertinib or a combination of Osimertinib and Tipifarnib. g: Overall survival of EGFR-L858R-T790M PDX mice treated with Osimertinib or the combination of Osimertinib and Tipifarnib. The graph is the result of one cohort of mice with n = 6 mice in both arms. P-value was determined by log-rank Mantel-Cox test. h: Representative images of Hematoxylin and Eosin (H&E) stainings (top) and Ki67 IHC (bottom) from PDX tumors collected after 2 weeks, 2 months and 5 months of treatment with Tipifarnib, Osimertinib or the combination of Osimertinib and Tipifarnib, respectively. Quantification of Ki67 IHC scores for each treatment. Data are representative of 3 to 6 independent tumors. Scale bar = 50μm i: Immunoblot of lysates from individual PDX harvested after 2 weeks, 2 months and 5 months of treatment with Tipifarnib, Osimertinib or the combination of Osimertinib and Tipifarnib, respectively. Quantification of protein levels of HRAS, p27Kip1 and phospho-Rb normalized with actin level. Images represent n = 3-4 independent tumors. Immunoblot was performed once. j: Proportion of G1, S and G2/M populations determined by scRNAseq Seurat clustering in PDX tumors collected after 2 weeks, 2 months and 5 months of treatment with Tipifarnib, Osimertinib or the combination of Osimertinib and Tipifarnib, respectively. k: GSEA analysis of the E2F_target signature in Osimertinib+Tipifarnib versus Osimertinib-treated EGFR-L858R-T790M PDX in the Ki67-negative (top) or positive (bottom) cell populations. l: Mean mRNA expression levels of genes implicated in replication or (m) in ISR, in the different cell populations, determined by scRNAseq. Mean ± SD are shown.

Here, we report that adaptive response to osimertinib is a highly dynamic process which invariably involves differentiation through an alveolar type-1 phenotype characteristically expressing cytoskeletal proteins involved in cellular contraction. Using a panel of Rho/ROCK pathway inhibitors, we found that tipifarnib, a clinically active farnesyltransferase inhibitor, efficiently and durably prevented relapse to osimertinib *in vitro* and *in vivo* by inducing an ATF4-dependent apoptotic response, with no evidence of toxicity in mice. Initially designed to inhibit RAS-driven cancer, FTIs did not achieve the expected clinical success, due to the lack of clarity of the clinical indication and the farnesylated proteins targeted for their pharmacological effects. Here we propose tipifarnib prevents the appearance of DTC and limit resistance to EGFR TKIs in EGFR-mutated lung cancer. In addition, we propose that concomitant inhibition of at least two farnesylated proteins RHOE and LaminB1 is sufficient to phenocopy tipifarnib’s effects suggesting that tipifarnib act on multiple targets. Our work opens the way to a new therapeutic combination to increase the efficacy of EGFR-targeted therapies.

## MATERIALS AND METHODS

### Cell culture

The human NSCLC cell-lines HCC4006 (CRL-2871, EGFR del L747-E749, A750P), HCC827 (CRL-2868, EGFR del E749-A750), HCC2935 (CRL-2869, EGFR del E746-T751, S752I) were obtained from the American Type Culture Collection (Manassas, VA, USA). The H3255 NSCLC cell line (EGFR L858R) and PC9 NSCLC cell-line (EGFR del E746-A750) were a kind gift from Helene Blons (APHP, Paris, France) and Antonio Maraver (IRCM, Montpellier, France) respectively. and were a kind gift from. For each cell-line, subclones were generated by limiting dilution. Moreover, subcultures were conducted for a limited period of time in order to limit cell deviation.

NSCLC cell-lines were cultured in RPMI (Roswell Park Memorial Institute) 1640 medium containing 10% fetal bovine serum (FBS) and were maintained at 37°C in a humidified chamber containing 5% CO2. Cell lines were authenticated and tested for mycoplasma contamination within the experimental time frame.

Cells were transduced with the Incucyte® Cell Cycle Green/Red lentivirus based on the FUCCI system, and selected by puromycin treatment for one week.

### SiRNA targeting RHOB, RHOE, LaminB1

Twenty-four hours after seeding, cells were transfected for 4h with siRNA (SMARTpool Dharmacon™) at a final concentration of 10nM, and 1µM osimertinib was added 24h hours after transfection. Transfection was repeated twice a week until relapse while maintaining osimertinib treatment. For combo transfection with two siRNAs, each siRNA was transfected at a final concentration of 5 nM and response to osimertinib was compared to cells transfected with siRandom at a final concentration of 10 nM.

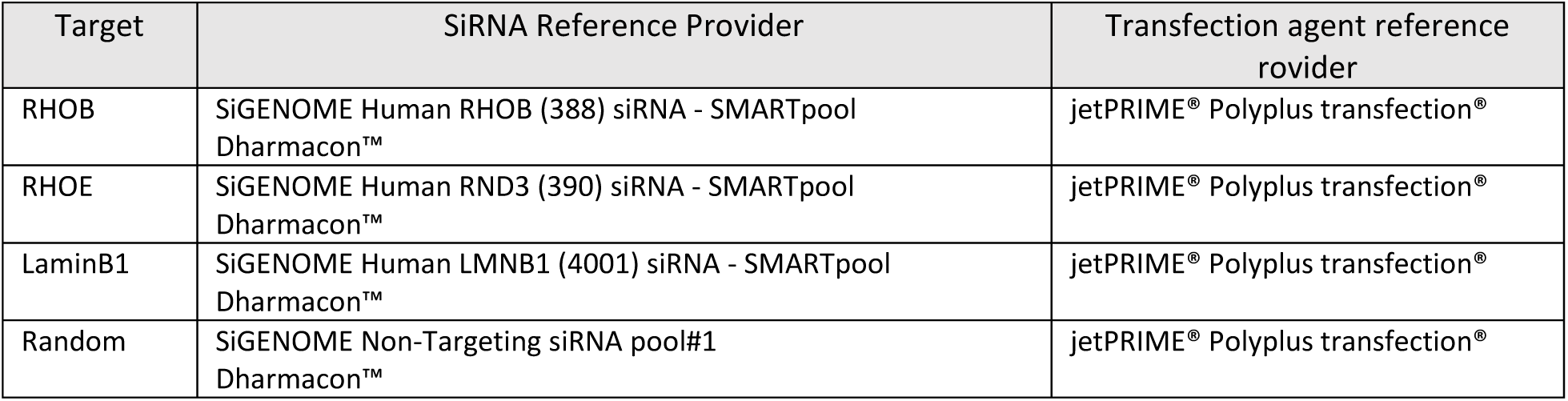

### Inhibitors

Cells were treated with the following inhibitors at the indicated concentrations two to three days after seeding, with medium changes twice a week for the whole duration of the experiment.

| Target | Drug | Provider | Reference | Concentrations used for *in vitro* experiments |
| --- | --- | --- | --- | --- |
| EGFR | Erlotinib (OSI-744, CP-358774, NSC 718781) | LC Laboratories | E4997 | 1µM |
|  | Osimertinib (AZD9291) | LC Laboratories | 0-7200 | 1µM |
| Farnesyl-transferase | Tipifarnib (R115777, IND 58359) | Kura Oncology |  | 0.1 - 1µM |
|  | Lonafarnib | Selleckchem | S2797 | 0.1 - 1µM |
|  | CP-609754 | Kura Oncology |  | 0.1 - 1µM |
|  | FTi-2153 | Kindly provided by Saïd Sebti and Andrew D. Hamilton |  | 1μM |
| Géranylgeranyl-transferase I | GGTI-298 | Selleckchem | S7466 | 1μM |
|  | GGTI-2166 | Kindly provided by Saïd Sebti and Andrew D. Hamilton |  | 1μM |
| ROCK | Y27632 | Selleckchem | S1049 | 10μM |
|  | GSK269962A | Selleckchem | S7687 | 0.3μM |
|  | RKI-1447 | Selleckchem | S7195 | 1μM |
| RHOA, B, C | TAT-C3 | C3 exoenzyme coupled to permeant peptide TAT (1 or 10 μg/ml) were produced and purified in our laboratory using an Akta purifier (GE Healthcare) as previously described (réf: Sahai, E. & Olson, M. F. Purification of TAT-C3 exoenzyme. Meth. Enzymol. 406, 128–140 (2006)). |  | 2 à 5 μg/ml |
| Actin | Latrunculin B | Sigma-Aldrich | 428020 | 0.3μM |
| CDK4/6 | Palbociclib | Selleckchem | S1579 | 1μM |

### Cell growth and viability assays on GFP-positives cells

Cells were transduced with a lentiviral vector (pLenti CMV GFP DEST (736-1), 9216bp) to obtain a constitutive expression of GFP protein. GFP-positives cells were selected by cell sorting using FACS Melody (BD Biosciences). Fluorescence intensity in each well of 96-well plate was evaluated twice a week using Synergy™ 2 Multi-Detection Microplate Reader. Relative cell survival in the presence of inhibitors which is proportional to fluorescence intensity was normalized to untreated cells after background corrections. Images of GFP-positive cells were obtained with Incucyte® S3 Live-Cell Analysis software (Sartorius).

### Crystal violet staining

Untreated control cells and treated cells in culture plates were wash with PBS, fixed with paraformaldehyde 4 % solution for 10 min and stained with a solution containing PBS - 0.5% crystal violet (Sigma-Aldrich/Merck, ref: C3886) – 25% methanol for 10 minutes. After 3 washes and drying, cell staining was imaged with a ChemiDoc™ MP Imaging system (Bio-Rad).

### Cytotoxicity assay

Viability of untreated cells or treated cells for 5 days was assessed using CellTiter 96® AQueous One Solution Cell Proliferation Assay (MTS) (Promega®, ref:G3580) according to manufacturer’s instructions.

### Western blot analyses

For *in vitro* experiments, cells were lysed with RIPA buffer complemented with proteases- and phosphatases-inhibitors. For animal experiments, frozen tumors were ground and lysed with Tris-SDS 1% buffer complemented with proteases- and phosphatases-inhibitors. After sonication, protein content was quantified using Bradford method. Protein extracts were separated on SDS-PAGE and electrotransferred on polyvinylidene difluoride membranes. Blots were probed with primary antibodies reported in the following table:

| Target | Provider | Reference |
| --- | --- | --- |
| RhoB | Santa Cruz Biotechnology | Sc-180 |
| RhoA | Santa Cruz Biotechnology | Sc-418 |
| RhoC | Cell Signaling | #3430 |
| RhoE | Cell Signaling | #3664 |
| p-ERK1/2 Thr202/Tyr204 | Cell Signaling | #4377 |
| ERK | Santa Cruz Biotechnology | Sc-93 |
| p-EGFR Tyr1068 | Cell Signaling | #2234 |
| EGFR | Cell Signaling | #4267 |
| Vimentin | Cell Signaling | #3932 |
| E-cadherin | Cell Signaling | #3195 |
| N-cadherin | Cell Signaling | #4061 |
| SLUG | Cell Signaling | #9585 |
| SNAIL | Cell Signaling | #3879 |
| p-RB Ser807/811 | Cell Signaling | #9308 |
| RB | Cell Signaling | #9309 |
| p27 | Cell Signaling | #3686 |
| p53 | Cell Signaling | #48818 |
| Cyclin D1 | Cell Signaling | #2978 |
| Cyclin E1 | Cell Signaling | #20808 |
| p-MLC2 Ser19 | Cell Signaling | #3671 |
| MLC2 | Cell Signaling | #8505 |
| p-Cofilin Ser3 | Cell Signaling | #3313 |
| Cofilin | Cell Signaling | #5175 |
| PARP | Cell Signaling | #9542 |
| Caspase-3 cleaved | Cell Signaling | #9661 |
| Caspase-3 | Cell Signaling | #9662 |
| Lamin B1 | Abcam | Ab16048 |
| Actin | Merck Millipore | MAB1501 |

Detection was performed using peroxydase-conjugated secondary antibodies and chemiluminescent detection kit (Clarity™ Western ECL, Bio-Rad) with a ChemiDoc™ MP Imaging system (Bio-Rad).

### Immunoprecipitation

Cells protein lysates were immunoprecipitated with a specific antibody for the GTP-bound active form of RHOA, RHOB and RHOC GTPases (RH12) designed in the laboratory ^25,26^. Then RHOA-, RHOB- and RHOC-active form expression was revealed by western blot.

### Phase contrast microscopy

Cells were imaged with ZEISS Axio Vert.A1 microscope and analysed with ZEN software (ZEISS).

### Phalloidin staining

Cells were first fixed with paraformaldehyde 4 % solution, permeabilized with a PBS-BSA 0.1 %-Triton X-100 0.5 % solution, blocked with a PBS-BSA 1 %-Triton X-100 0.1 % solution and stained using Alexa Fluor™ 594 phalloidin (ThermoFisher Scientific, #A12381) diluted 50 times in a PBS-BSA 0.1 %-Triton X-100 0.1 % solution. Then, nuclei were stained with DAPI (4’, 6-diamidino-2-phenylindole, dihydrochloride, ThermoFisherScientific, #D1306). Images were acquired with ZEISS Axio Vert.A1 microscope and analysed with ZEN software (ZEISS). Quantification of F-actin intensity was performed by ImageJ software

### Cell cycle analysis

Cells were transduced with Incucyte® Cell Cycle Green/Red Lentivirus Reagent (EF1α-Puro) (Sartorius, Cat. No. 4779) as recommended by the manufacturer and transduced cells were selected by puromycin treatment. Cells were imaged every hour with Incucyte® S3 Live-Cell Analysis System (Sartorius). G1-phase, S/G2/M-phase and G1/S-phase cells were quantified using Incucyte® S3 Live-Cell Analysis software (Sartorius).

### RNA sequencing

Subclones generated from HCC4006, PC9 and H3255 cell lines were treated with 1µM erlotinib for 24h, 21 days (DTC state) and cultured for more than 3 months in the presence of the drug (RPC state, Resistant Proliferative Clones) for HCC4006 and PC9. HCC827 were treated until reaching the drug-tolerant state (11 days) with 1 µM osimertinib. RNA extraction was performed using AllPrep DNA/RNA Mini kit (Qiagen, #80204) according to manufacturer’s protocol. RNA quality was assessed using Fragment Analyzer (Agilent technologies) and the RQN values were provided to confirm the integrity of total RNA. RNA concentration was determined by fluorescent method using Quant-iT™ RNA Assay Kit, Broad Range (ThermoFisher Scientific). RNA samples were processed with Illumina TruSeq® Stranded mRNA Library Preparation Kit following the manufacturer’s protocol. Library size and quality were confirmed on Fragment analyzer (Agilent Technologies). KAPA quantification kit for Illumina platforms (KAPA Biosystems, Roche) was used to quantify library by qPCR. Indexed libraries were pooled and sequenced on an Illumina NextSeq 550 (2×75 bp paired-end reads).

Reads were mapped and counted using the RNA-Seq by Expectation Maximization (RSEM) software v. 1.3 (with bowtie2-2.3.5.1) based on the human reference genome UCSC hg38. Differential expression was analysed with DESeq2.

### Single-cell capture

#### HCC4006-treated cells

HCC4006 subcloned cells were expanded in untreated, osimertinib- or osimertinib + tipifarnib-containing medium for 4, 20 and 12 days respectively. Medium was changed 24h before sorting. Cells were dissociated by trypsinisation, recovered in FACS buffer (0,04% BSA in PBS) and kept on ice. G1 (red) and S/G2 (green) cells were sorted at 4°C using FACS Melody (BD Biosciences). Cells were spun down and resuspended in appropriate volume for a final concentration of 500 to 1500 cells / µl, with a viability above 85%. 6000 cells per sample were loaded to the Chromium Controller (10X Genomics) and 1900-3300 cells were recovered depending on the sample. scRNA-seq libraries were generated using the 10X Genomics Chromium Single Cell 3’ Kit v2 according to manufacturer’s instructions. The libraries were profiled with the HS NGS kit for the Fragment Analyzer (Agilent Technologies) and quantified using the KAPA library quantification kit (Roche Diagnostics). The libraries were pooled and sequenced on the Illumina NextSeq550 instrument by the CRCT Genomics platform using the High Output 150 cycles kit. The single-indexed sequencing parameters were 28, 8, 0, 91 cycles (read 1, index 1, index 2, read 2). The dual-indexed sequencing parameters were 28, 10, 10, 90 cycles (read 1, index 1, index 2, read 2). An average depth of ∼46000 reads/cell was obtained.

#### Patient-Derived Xenograft

The PDX model (EGFR^T790M/L858R^, TP103^21^) was generated in the Paz-Ares laboratory at the Instituto de Biomedicina de Sevilla (IBIS). Tumours were collected, cut into 2-4 mm pieces on ice and dissociated by mechanical and enzymatic reaction following the Miltenyi Tumour Dissociation Kit instructions (GentleMACS program h_tumor_02). Samples were filtered through a MACS SmartStrainer 70 µm. Erythrocytes were lysed following the Red Blood Cell Lysis Solution protocol (Miltenyi Biotec). Samples were frozen in RPMI 10% DMSO 20% FBS until the day of single-cell capture. Samples from all conditions were thawed and processed on the same day following the 10X Genomics Thawing Dissociated Tumor Cells for Single Cell RNA Sequencing protocol. Cells were recovered in appropriate volume for a final concentration of 500 to 1000 cells / µl, with a viability above 70%. 4500 cells per sample were loaded to the Chromium Controller (10X Genomics) and scRNA-seq libraries were generated using the 10X Genomics Chromium Single Cell 3’ Kit v2 according to manufacturer’s instructions. The libraries were sequenced with a NextSeq 550 (Illumina) by the CRCT Genomics platform. An average depth of ∼27000 reads/cell was obtained.

### Single cell RNA sequencing analysis

The single cell transcriptomic data was loaded from Kurppa et al.^4^ on GEO under accession number GSE131604. The data from the different conditions were pulled together to create a global counts matrix. A dimension reduction by principal component analysis (PCA), a t-SNE projection and a k-means clustering were done using the clustering function of the SingleCellSignalR R package. Two differential gene expression analyses were done using the cluster_analysis function of SingleCellSignalR using respectively the cluster vector from the clustering function cited above and a cluster vector created to correspond to the different conditions. The p-value threshold was set by default at 5%. A pathway signature genes table was retrieved from Reactome and GO. The mean expression of the genes in each pathway was calculated in each cell to create a mean pathway expression matrix. Violin plots of the mean pathway expression were computed and displayed using the ggplot2 package. The heatmaps were generated using the ComplexHeatmap R package. The t-SNE expression plots were done using the expression_plot function of SingleCellSignalR. All the analyses were carried out with RStudio.

### Mouse xenograft studies

Cell line xenograft experiments were performed in 6 to 8-week old female NMRI mice (Charles River Laboratories) by injecting 10 million PC9 cells in 50% matrigel subcutaneously on each flank of the mouse. The PDX model (EGFR^T790M/L858R^, TP103^21^) was cut and engrafted subcutaneously into the flank of NSG mice.

When tumors reached on average 200 to 300 mm^3^ (calculated as [length x width^2^ x 3.14/6]), mice were randomized and treated by oral gavage 5 days a week with 100µl of vehicle, osimertinib (5 mg/kg q.d), tipifarnib (80 mg/kg b.i.d) or a combination of osimertinib and tipifarnib. Osimertinib was diluted in 0,5% carboxymethylcellulose (Sigma, M0512) + 0,1% Tween80 (Sigma, P8074) and Tipifarnib in 2-hydroxypropryl-cyclodextrin 20% (Sigma, H107). Tumors were measured twice a week. Mice were weighted twice a week to monitor toxicity and humanely killed at indicated times and tumors harvested.

### Immunohistochemistry

Tissues were fixed in 10% formalin overnight at room temperature, stored in 70% ethanol at 4°C and embedded in paraffin. Four-micrometer paraffin sections were used for hematoxylin and eosin staining followed by immunohistochemistry using standard procedures with Ki67 antibody (SP6; Thermo Scientific). Quantification of Ki67 IHC scores for each treatment was blind evaluated by two operators.

### TCGA and GSEA analysis

RSEM-normalized expression data of human lung adenocarcinomas (517 samples) and healthy lungs (59 samples) from The Cancer Genome Atlas (TCGA) database were downloaded from http://firebrowse.org. Spearman’s correlation and statistical analysis of mRNA expression in LUAD was performed using the cbioportal co-expression tool (www.cbioportal.org). Pathway Gene Set Enrichment Analysis (GSEA) was performed with dedicated software (https://www.gsea-msigdb.org/gsea).

## ACKNOWLEDGMENTS

We thank all members of the SIGNATHER team at CRCT (Cancer Research Center of Toulouse, INSERM U1037, CNRS ERL5294, UPS, Toulouse, France) for stimulating and thoughtful discussions. We thank David Santamaría, John Hickman, Agnese Cristini and Olivier Sordet for critically reading the manuscript. We thank all members of mice core facilities (CREFRE, UMS006 Inserm, Toulouse) in particular Cédric Baudelin, Amanda Corini, Emilie Sinhlivong, Tristan Chapon, Charlotte Delos, Thi-Lan Huong Huynh and Charlène Lopez for their support and technical assistance. We acknowledge the cytometry/cell-sorting and imaging facilities of the CRCT, in particular Manon Farcé and Laetitia Ligat for their assistance with flow cytometry and microscopy, respectively. We thank Carine Valle and Emeline Sarot from the Genomics and Transcritomics platform of the CRCT for their support and technical assistance, as well as Marie Tosolini from the Bioinformatics platform of the CRCT for the bioinformatics analysis. We thank Géraldine Touriol for the administrative and financial management.

## AUTHOR CONTRIBUTIONS

Study design, data interpretation and preparation of the manuscript: SF, CD, RG, AP, JM, OC, GF; Execution of experiments: SF, CD, RG, RA, SP, CT, EC, AC, OC; Computational and statistical analysis: JC, JPV; Writing, review, and/or revision of the manuscript: SF, CD, RG, JPV, AM, AP, JM, OC, GF; Material support: ILM, IF, LPA.

## FUNDINGS

This work was supported in part by institutional grants from INSERM, Fondation pour la Recherche Médicale (FRM, équipe labellisée [DEQ20170839117]), Programme de Recherche Translationnelle en Cancérologie (INCa-DGOS, [PRT-K18-048], JM), Fondation Toulouse Cancer Santé (FTCS, OC and JM), Labex TOUCAN (OC), Ligue Nationale Contre le Cancer (LNCC, GF).

## CONFLICT OF INTEREST STATEMENT

Part of this study received financial support from Kura Oncology through a sponsored research contract.

## DATA AVAILABILITY

RNAseq and scRNAseq raw data will be available upon request through private access and publicly available upon acceptance of the publication.

## Notes

### Competing Interest Statement

The authors have declared no competing interest.

## REFERENCES

1 Sharma, S. V. et al. A chromatin-mediated reversible drug-tolerant state in cancer cell subpopulations. Cell 141, 69–80, doi:10.1016/j.cell.2010.02.027 (2010).

2 Hata, A. N. et al. Tumor cells can follow distinct evolutionary paths to become resistant to epidermal growth factor receptor inhibition. Nature medicine 22, 262–269, doi:10.1038/nm.4040 (2016).

3 Ramirez, M. et al. Diverse drug-resistance mechanisms can emerge from drug-tolerant cancer persister cells. Nature communications 7, 10690, doi:10.1038/ncomms10690 (2016).

4 Kurppa, K. J. et al. Treatment-Induced Tumor Dormancy through YAP-Mediated Transcriptional Reprogramming of the Apoptotic Pathway. Cancer cell 37, 104–122 e112, doi:10.1016/j.ccell.2019.12.006 (2020).

5 Guler, G. D. et al. Repression of Stress-Induced LINE-1 Expression Protects Cancer Cell Subpopulations from Lethal Drug Exposure. Cancer cell 32, 221–237 e213, doi:10.1016/j.ccell.2017.07.002 (2017).

6 Vallette, F. M. et al. Dormant, quiescent, tolerant and persister cells: Four synonyms for the same target in cancer. Biochem Pharmacol 162, 169–176, doi:10.1016/j.bcp.2018.11.004 (2019).

7 Sakaue-Sawano, A. et al. Visualizing spatiotemporal dynamics of multicellular cell-cycle progression. Cell 132, 487–498, doi:10.1016/j.cell.2007.12.033 (2008).

8 Turke, A. B. et al. Preexistence and clonal selection of MET amplification in EGFR mutant NSCLC. Cancer cell 17, 77–88, doi:10.1016/j.ccr.2009.11.022 (2010).

9 Oren, Y. et al. Cycling cancer persister cells arise from lineages with distinct programs. Nature 596, 576–582, doi:10.1038/s41586-021-03796-6 (2021).

10 Maynard, A. et al. Therapy-Induced Evolution of Human Lung Cancer Revealed by Single-Cell RNA Sequencing. Cell 182, 1232–1251 e1222, doi:10.1016/j.cell.2020.07.017 (2020).

11 Travaglini, K. J. et al. A molecular cell atlas of the human lung from single-cell RNA sequencing. Nature 587, 619–625, doi:10.1038/s41586-020-2922-4 (2020).

12 Rottner, K., Faix, J., Bogdan, S., Linder, S. & Kerkhoff, E. Actin assembly mechanisms at a glance. Journal of cell science 130, 3427–3435, doi:10.1242/jcs.206433 (2017).

13 Sit, S. T. & Manser, E. Rho GTPases and their role in organizing the actin cytoskeleton. Journal of cell science 124, 679–683, doi:10.1242/jcs.064964 (2011).

14 Calvayrac, O. et al. The RAS-related GTPase RHOB confers resistance to EGFR-tyrosine kinase inhibitors in non-small-cell lung cancer via an AKT-dependent mechanism. EMBO molecular medicine 9, 238–250, doi:10.15252/emmm.201606646 (2017).

15 Delmas, A. et al. The c-Jun/RHOB/AKT pathway confers resistance of BRAF-mutant melanoma cells to MAPK inhibitors. Oncotarget 6, 15250–15264, doi:10.18632/oncotarget.3888 (2015).

16 Shu, Z., Row, S. & Deng, W. M. Endoreplication: The Good, the Bad, and the Ugly. Trends Cell Biol 28, 465–474, doi:10.1016/j.tcb.2018.02.006 (2018).

17 Gandarillas, A., Molinuevo, R. & Sanz-Gomez, N. Mammalian endoreplication emerges to reveal a potential developmental timer. Cell Death Differ 25, 471–476, doi:10.1038/s41418-017-0040-0 (2018).

18 Torrence, M. E. et al. The mTORC1-mediated activation of ATF4 promotes protein and glutathione synthesis downstream of growth signals. Elife 10, doi:10.7554/eLife.63326 (2021).

19 Pakos-Zebrucka, K. et al. The integrated stress response. EMBO Rep 17, 1374–1395, doi:10.15252/embr.201642195 (2016).

20 Storck, E. M. et al. Dual chemical probes enable quantitative system-wide analysis of protein prenylation and prenylation dynamics. Nat Chem 11, 552–561, doi:10.1038/s41557-019-0237-6 (2019).

21 Quintanal-Villalonga, A. et al. FGFR1 Cooperates with EGFR in Lung Cancer Oncogenesis, and Their Combined Inhibition Shows Improved Efficacy. Journal of thoracic oncology : official publication of the International Association for the Study of Lung Cancer 14, 641–655, doi:10.1016/j.jtho.2018.12.021 (2019).

