## Supplemental data for "Farnesyltransferase inhibition overcomes the adaptive resistance to osimertinib in *EGFR*-mutant NSCLC"

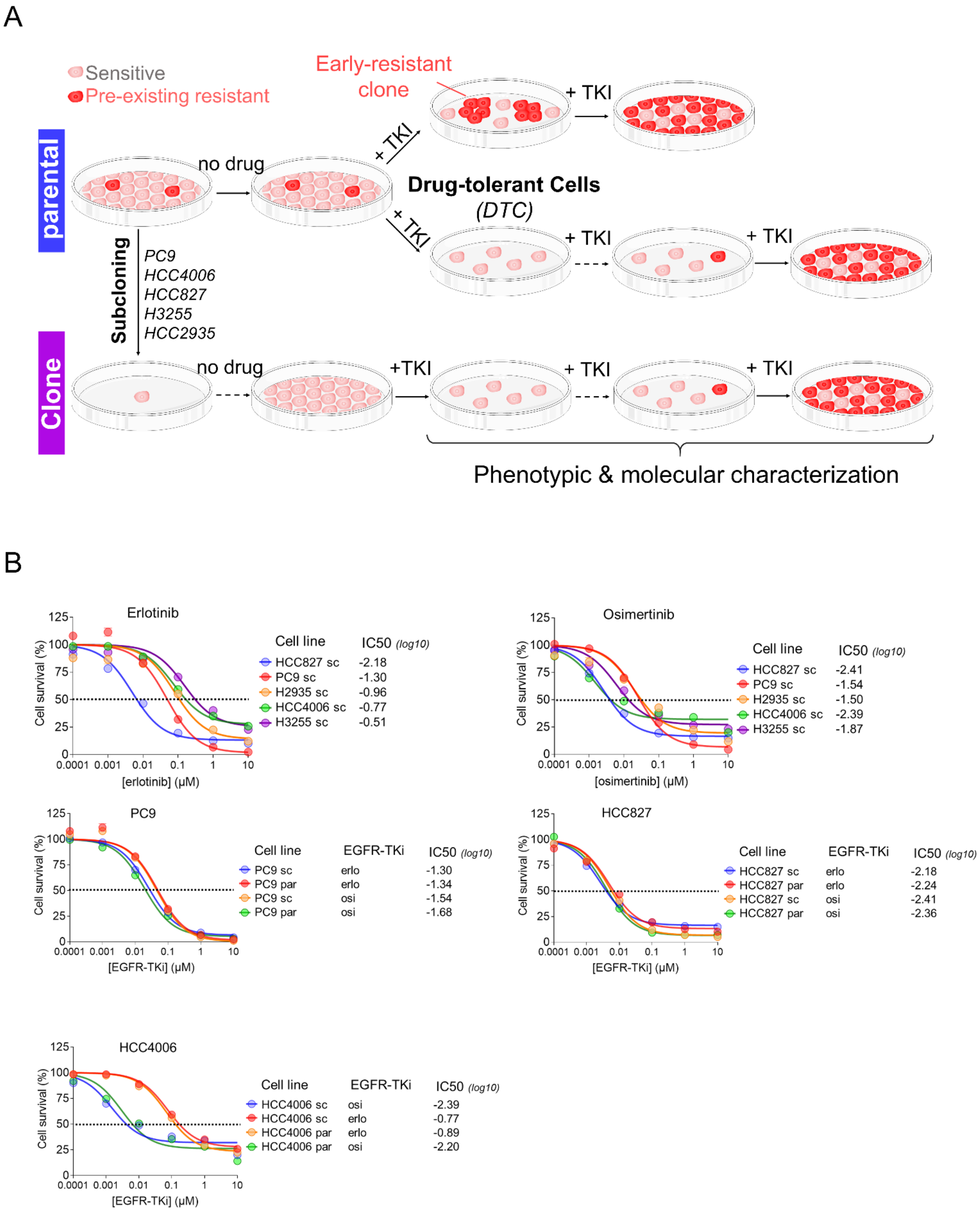


**Supp Figure 1:**

A: Experimental design. Parental EGFR-mutated PC9, HCC4006, HCC827, H3255 and HCC2935 cells were subcloned in order to avoid/minimize the presence of potential pre-existing EGFR-TKi-resistant cells. Then DTC and RPC were characterized at phenotypic and molecular level.

B: Cell survival (%) by cytotoxicity assay of parental (par) and subclonal (sc) PC9, HCC4006, HCC827, H3255 and HCC2935 cells upon erlotinib or osimertinib treatment for 5 days. Half inhibitory concentration for each condition were evaluated (log10)

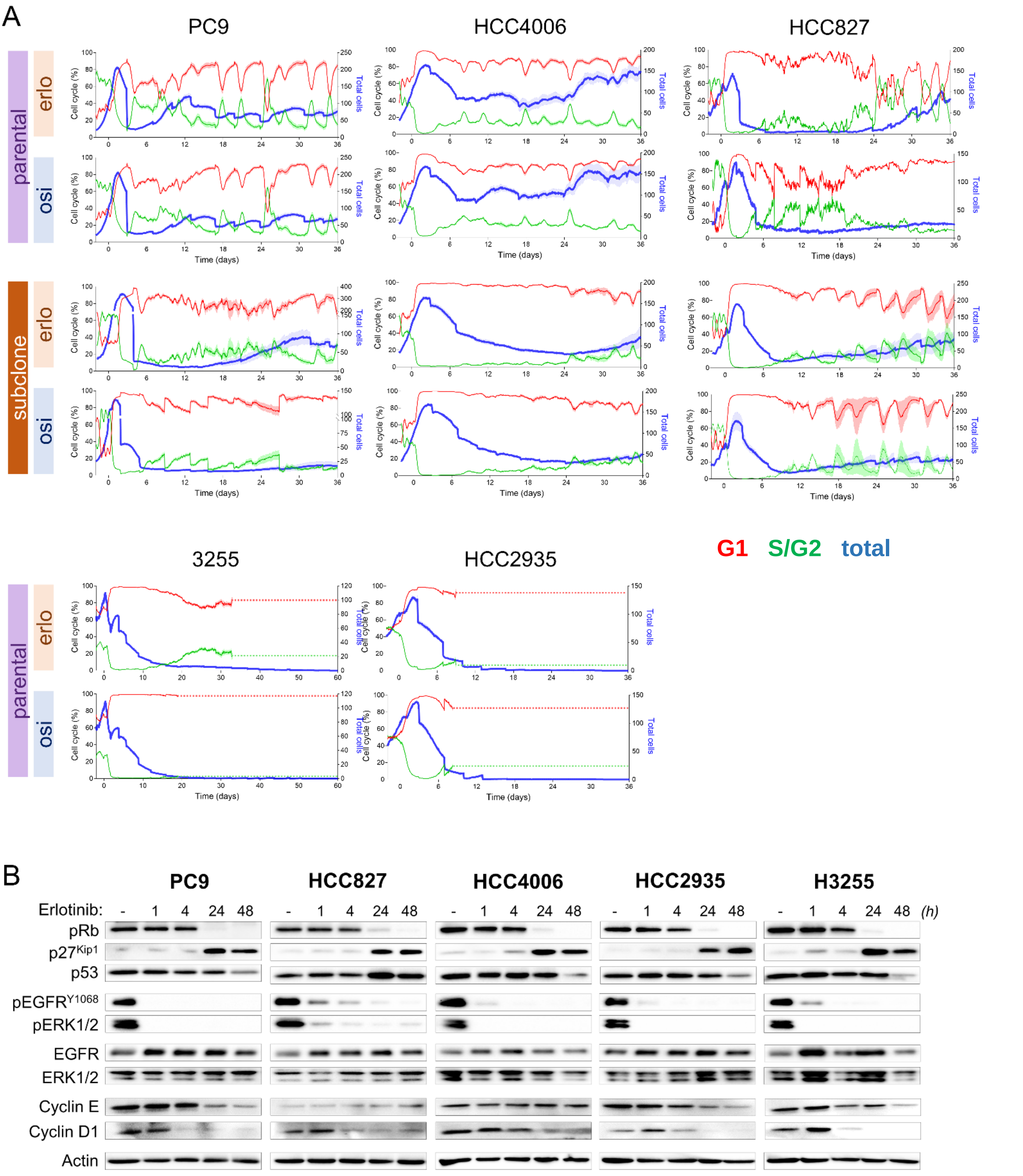


**Supp Figure 2:**

A: Cell cycle analysis of HCC4006, PC9, HCC2935, HCC827 parental (par) or subclonal (sc) cells treated with erlotinib or osimertinib (1µM). Top panels: Fraction of total parental or subclonal cells (%) according to day 0. Middle panels: Distribution of G1 (red), S/G2 (green) and G1-S (yellow) parental cells (%). Bottom panels: Distribution of G1 (red), S/G2 (green) and G1-S (yellow) sublconal cells (%). DTC state refers to the period between the onset of cell mortality and the recovery of cell proliferation.

B: Protein expression by western blot of proteins related to EGFR pathway (phospho-EGFR, EGFR, phospho-ERK, ERK) and cell cycle (p27Kip1, p53, phospho-RB, RB, cyclin E, cyclin D1) on PC9, HCC4006, HCC827, HCC2935 and H3255 cells treated with erlotinib (1µM).

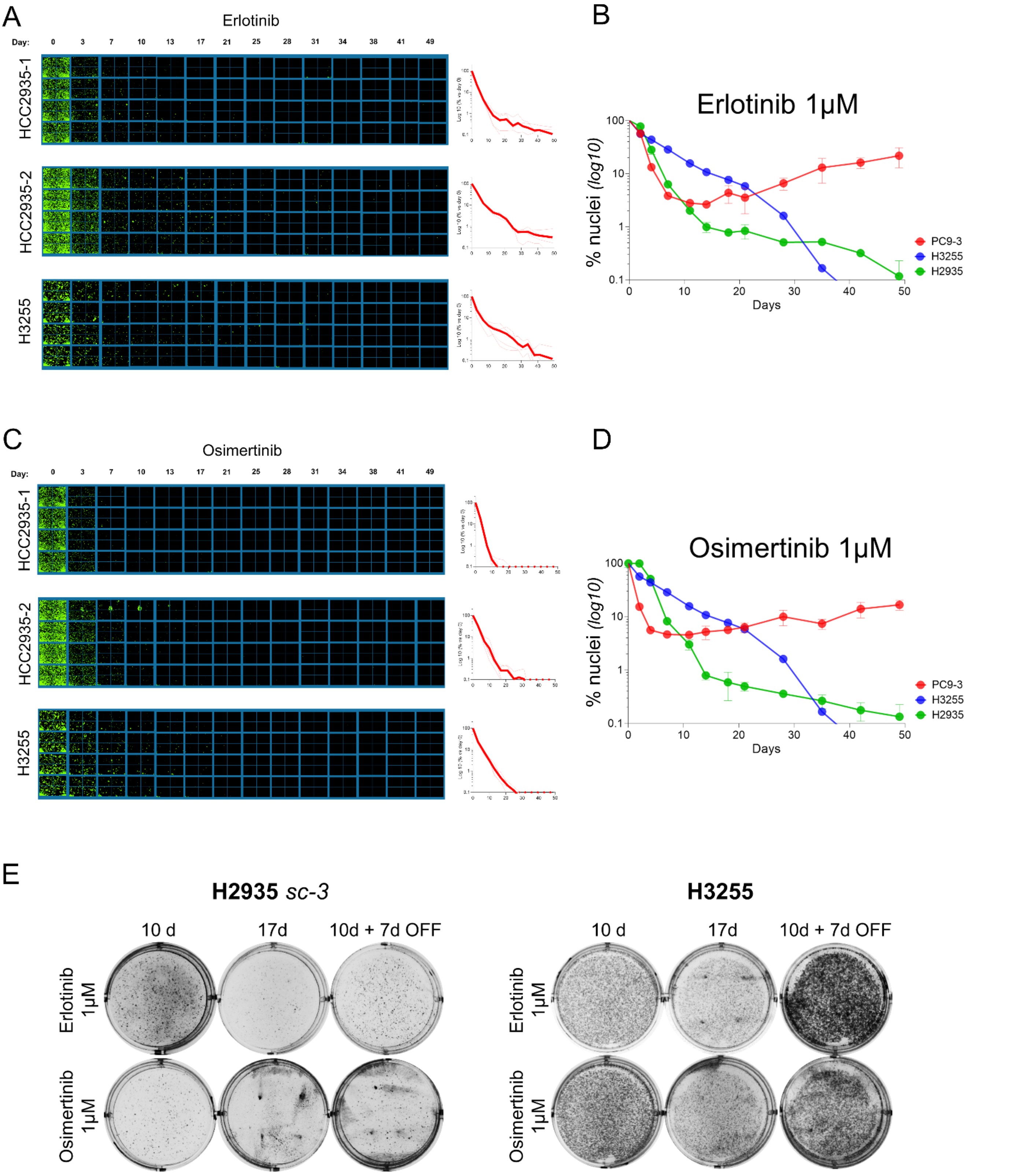


**Supp Figure 3:**
A: Images of GFP-positive HCC2935 subclonal and H3255 parental cells upon erlotinib (1µM) treatment. Graphs represent cell survival (log10 % vs day 0).
B: Cell survival (log10 % vs day 0) of PC9 subclonal, HCC2935, H3255 parental cell upon erlotinib (1µM) treatment.
C: Images of GFP-positive HCC2935 subclonal and H3255 parental cells upon osimertinib (1µM) treatment. Graphs represent cell survival (log10 % vs day 0).
D: Cell survival (log10 % vs day 0) of PC9 subclonal, HCC2935, H3255 parental cell upon osimertinib (1µM) treatment.
E: Cell survival by with crystal violet staining of HCC2935 subclonal and H3255 parental cells treated for 10 or 17 days with erlotinib or osimertinib (1µM) and at 17 days after 7 days of drug withdrawal.

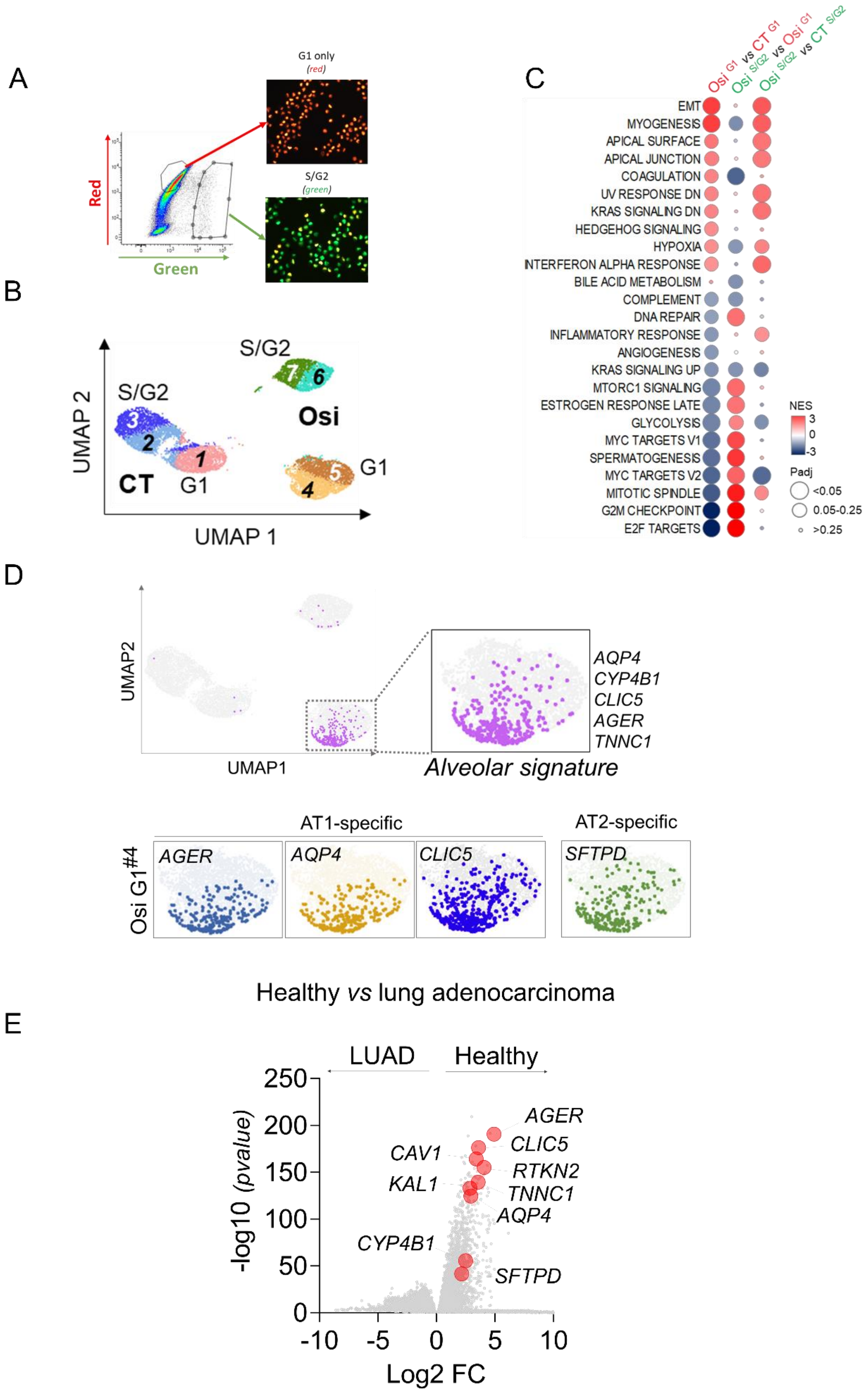


**Supp Figure 4:**
A: Flow cytometry charts of red cells (G1) or green cells (S/G2 or G1-S) of osimertinib-treated (1µM, 20 days) HCC4006 subclonal cells sorted for single-cell RNA sequencing experiment.
B: UMAP representation of the different population of untreated and osimertinib-treated HCC4006 clonal cells.
C: GSEA of Hallmark gene sets comparing Osimertinib-treated G1 versus untreated-G1 HCC4006 cells, Osimertinib-treated S/G2 versus Osimertinib-treated G1 HCC4006 cells or Osimertinib-treated S/G2 versus untreated S/G2 HCC4006 cells.
D: UMAP representation of the alveolar signature. Cells positive for AGER, AQP4, CLIC5 or SFTPD are shown.
E: Violin plot of the differentially expressed genes between healthy and lung adenocarcinomas (LUAD) using TCGA database.

Figarol *et al.*

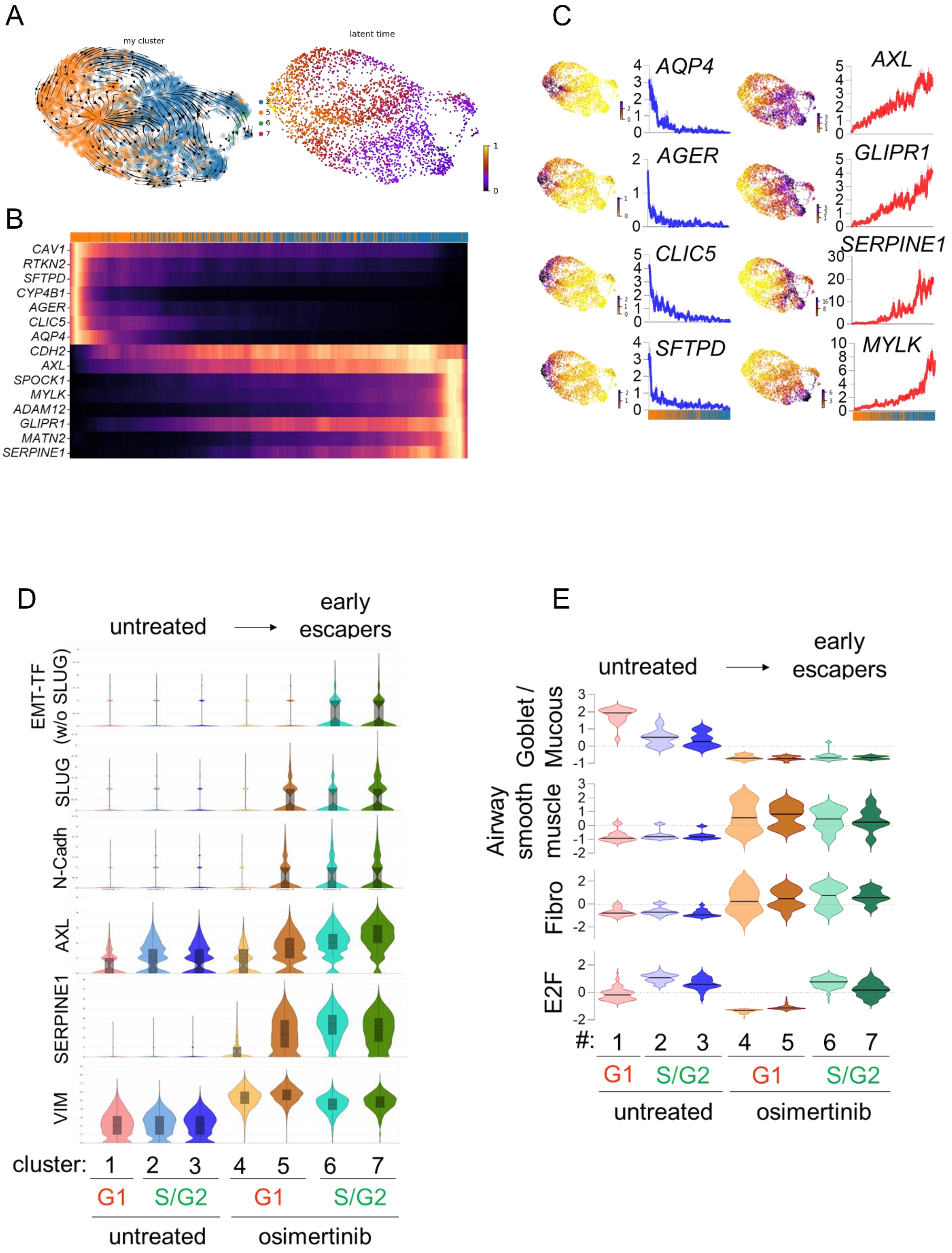


**Supp Figure 5:**

A: Velocity and clustering (left) and latent time (right) analysis of the osimertinib-treated G1 subpopulation.

B: Normalized mRNA expression of alveolar and EMT-related genes according to latent time in the osimertinib-treated G1 subpopulation.

C. Repartition of AT1- and EMT-related genes expression within the osimertinib-treated G1 cluster and z-score normalized mRNA expression of corresponding genes according to the latent time.

D: Violin plots representing the Log2 mRNA expression of EMT-related genes significantly overexpressed during osimertinib treatment in the different clusters.

E: Mean z-score of normal cell type-associated signatures (Maynard, Nature 2020) significantly regulated by osimertinib treatment in the different clusters.

Figarol *et al.*

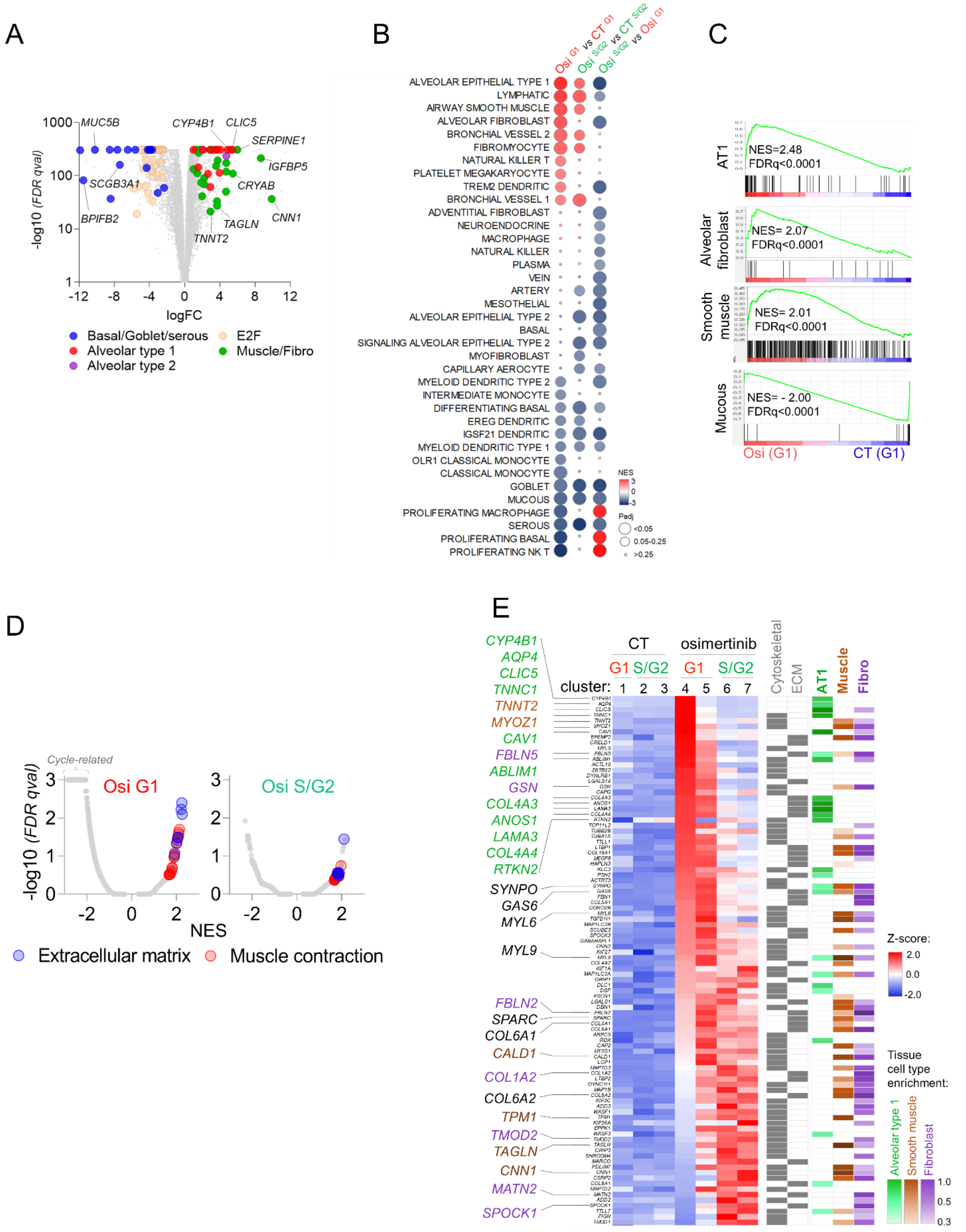


**Supp Figure 6:**

A: Volcano plot of the differentially expressed genes between osimertinib-G1 and untreated-G1 HCC4006 clonal cells

B: GSEA of normal lung cell types signatures (Maynard, Nature, 2020) comparing Osimertinib-treated G1 versus untreated-G1 HCC4006 cells, Osimertinib-treated S/G2 versus Osimertinib-treated G1 HCC4006 cells or Osimertinib-treated S/G2 versus untreated S/G2 HCC4006 cells.

C: GSEA analysis of significantly regulated normal lung cell types signatures between osimertinib-treated G1 versus untreated-G1 HCC4006 clonal cells.

D: Volcano plot of the differentially regulated C5 gene signatures revealed by GSEA analysis between osimertinib-G1 vs untreated-G1 or osimertinib-S/G2 vs untreated S/G2 HCC4006 clonal cells

E: Normalized mRNA expression (z-score) of cytoskeletal and extracellular matrix genes upregulated by osimertinib treatment (p<0.01, Log2FC>1). Tissue cell type enrichment is shown.

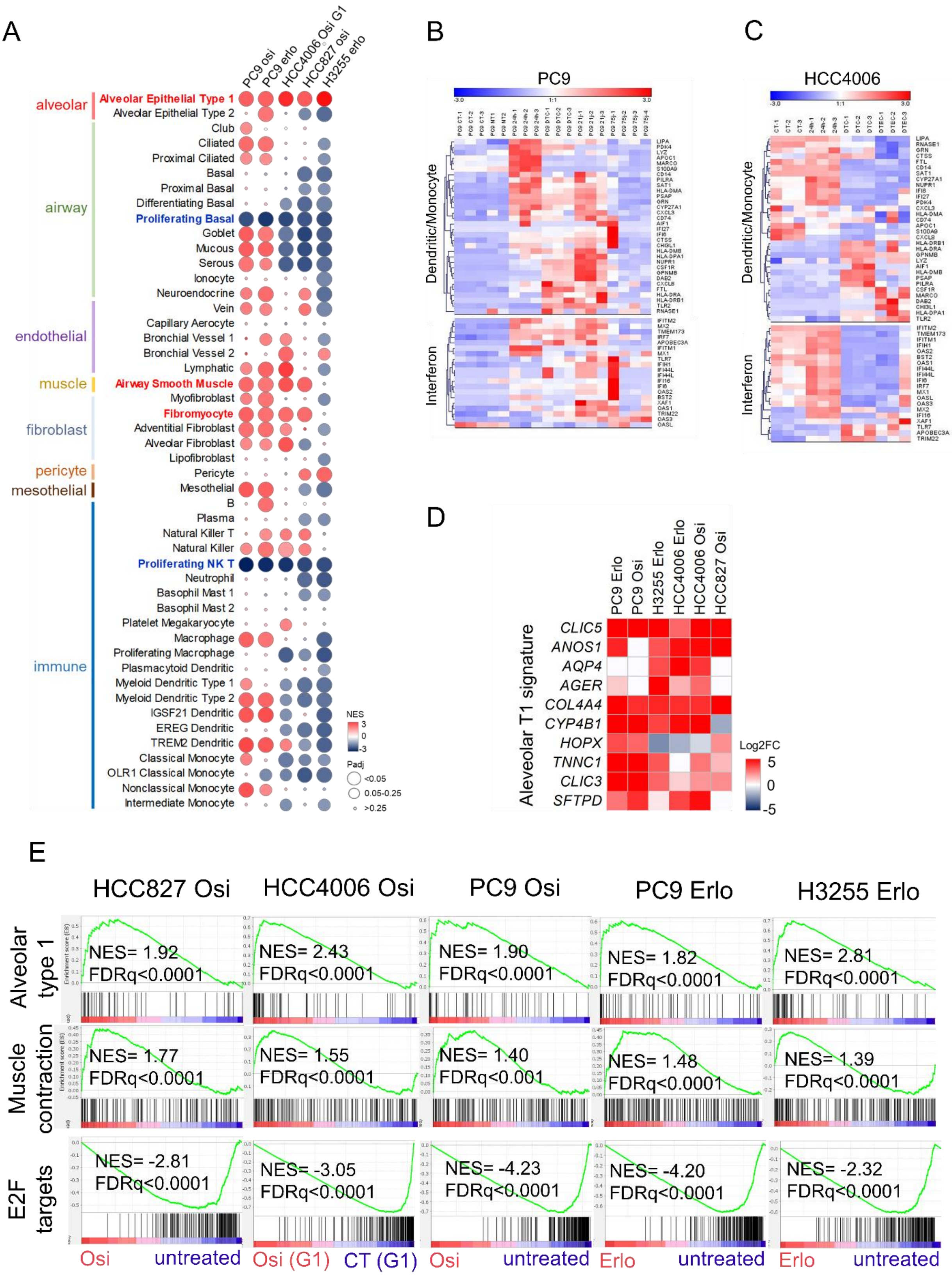


**Supp Figure 7:**

A: GSEA analysis of normal lung cell type signatures in erlotinib or osimertinib-derived DTC generated from the indicated cell lines

B: Normalized mRNA expression (z-score) of Dendritic/monocyte- (top) or interferon- (bottom) associated genes in erlotinib-treated PC9 clonal cells (24h, 7d, 21d, 75d).

C: Normalized mRNA expression (z-score) of Dendritic/monocyte- (top) or interferon- (bottom) associated genes in erlotinib-treated HCC4006 clonal cells (24h, 21d, 150d).

D: Differential mRNA expression of alveolar type 1-related genes in DTC generated from indicated cell lines and treatments, compared to corresponding untreated cells

E: GSEA analysis of alveolar type 1, muscle contraction and E2F_target signatures in DTC generated from indicated cell lines and treatments

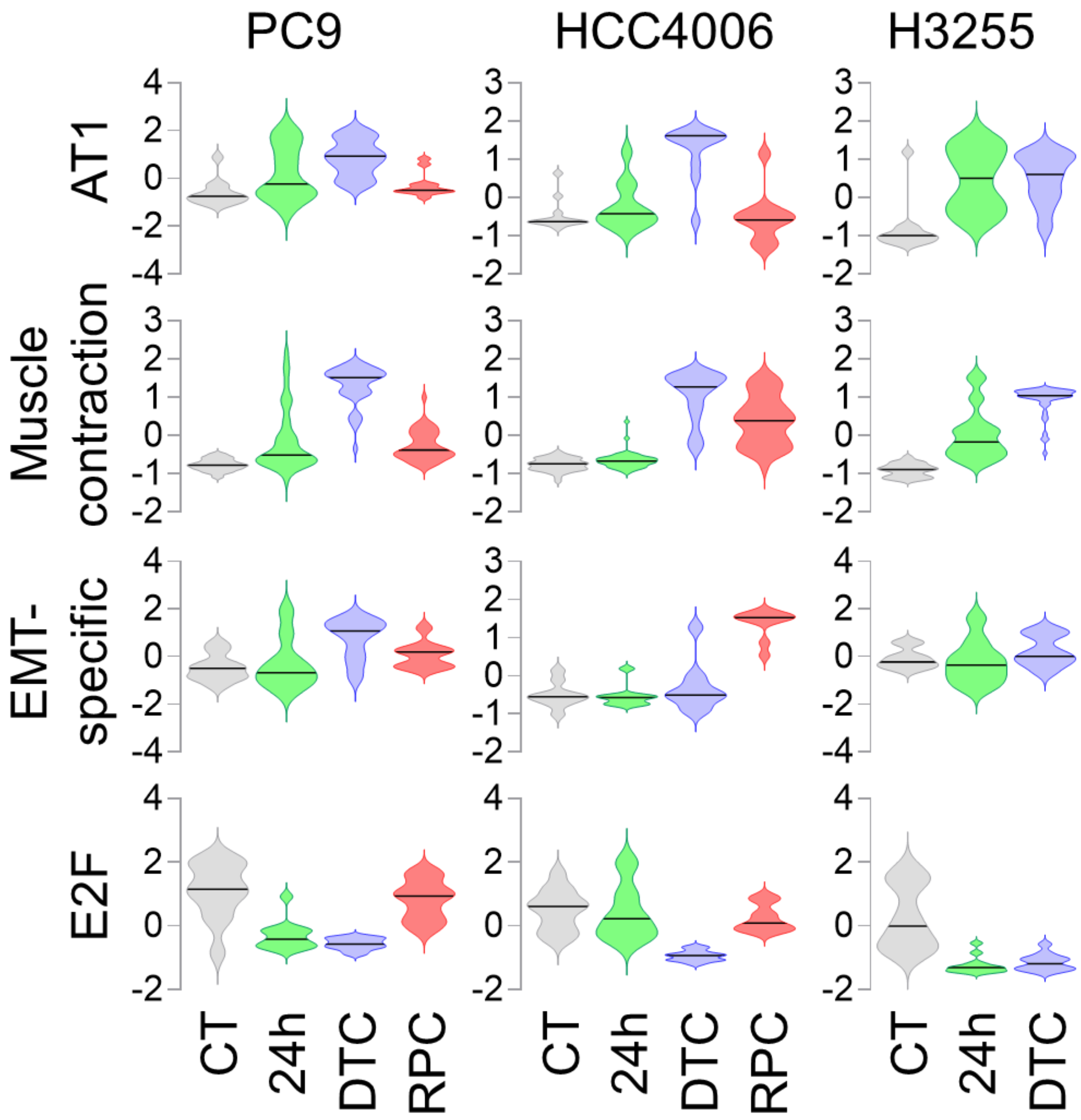


**Supp Figure 8:**
Violin plots showing the mean z-score evolution of alveolar type 1 (AT1), muscle contraction, EMT-specific and E2F_targets signatures in erlotinib-treated PC9, HCC4006, H3255 cells.

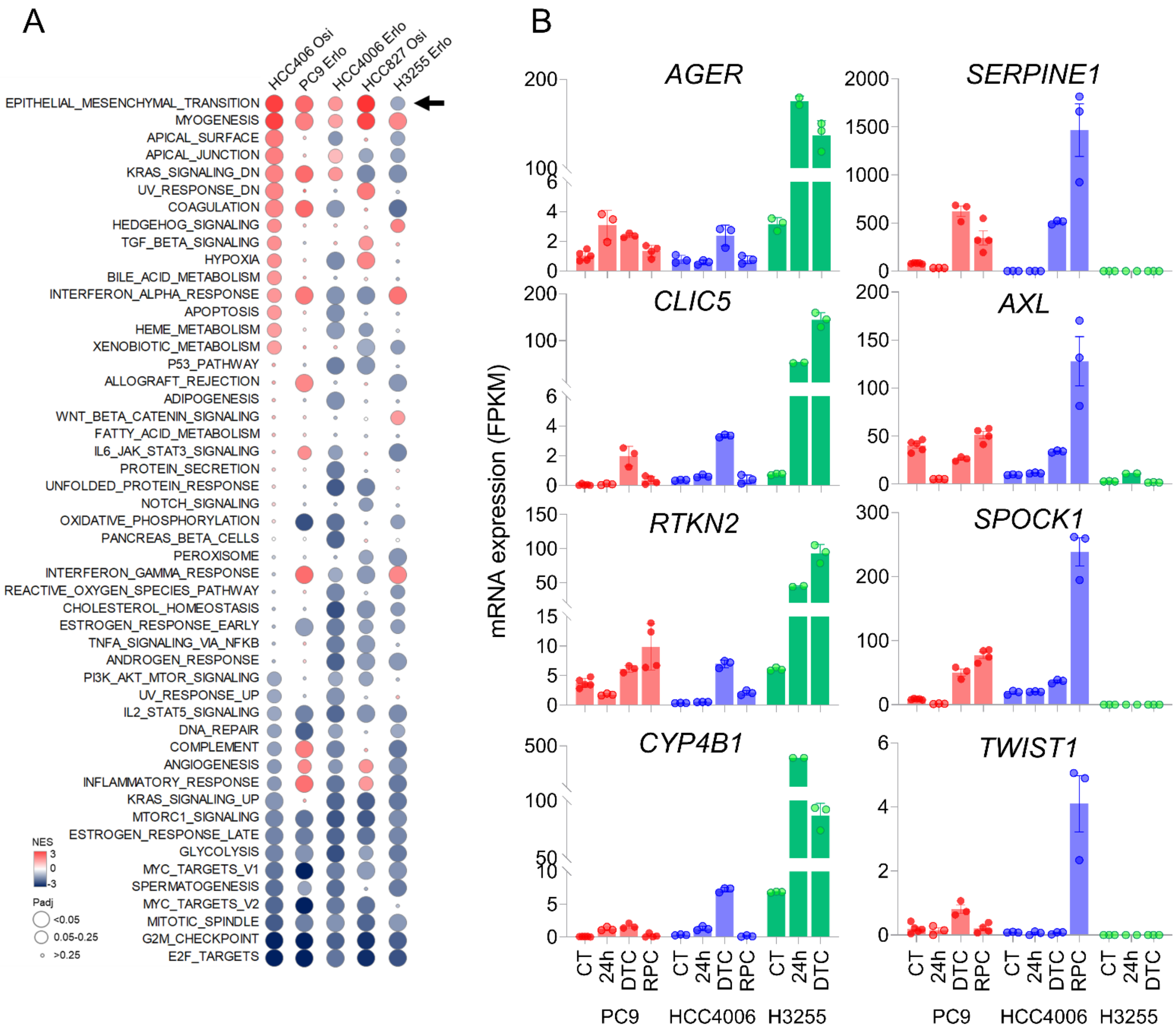


**Supp Figure 9:**
A: GSEA of Hallmark gene sets in DTC generated in indicated cell lines
B: mRNA expression levels of AT1- (left) or EMT- (right) associated genes in erlotinib-treated PC9, HCC4006 and H3255 cells. Mean FPKM ± SEM from RNAseq data.

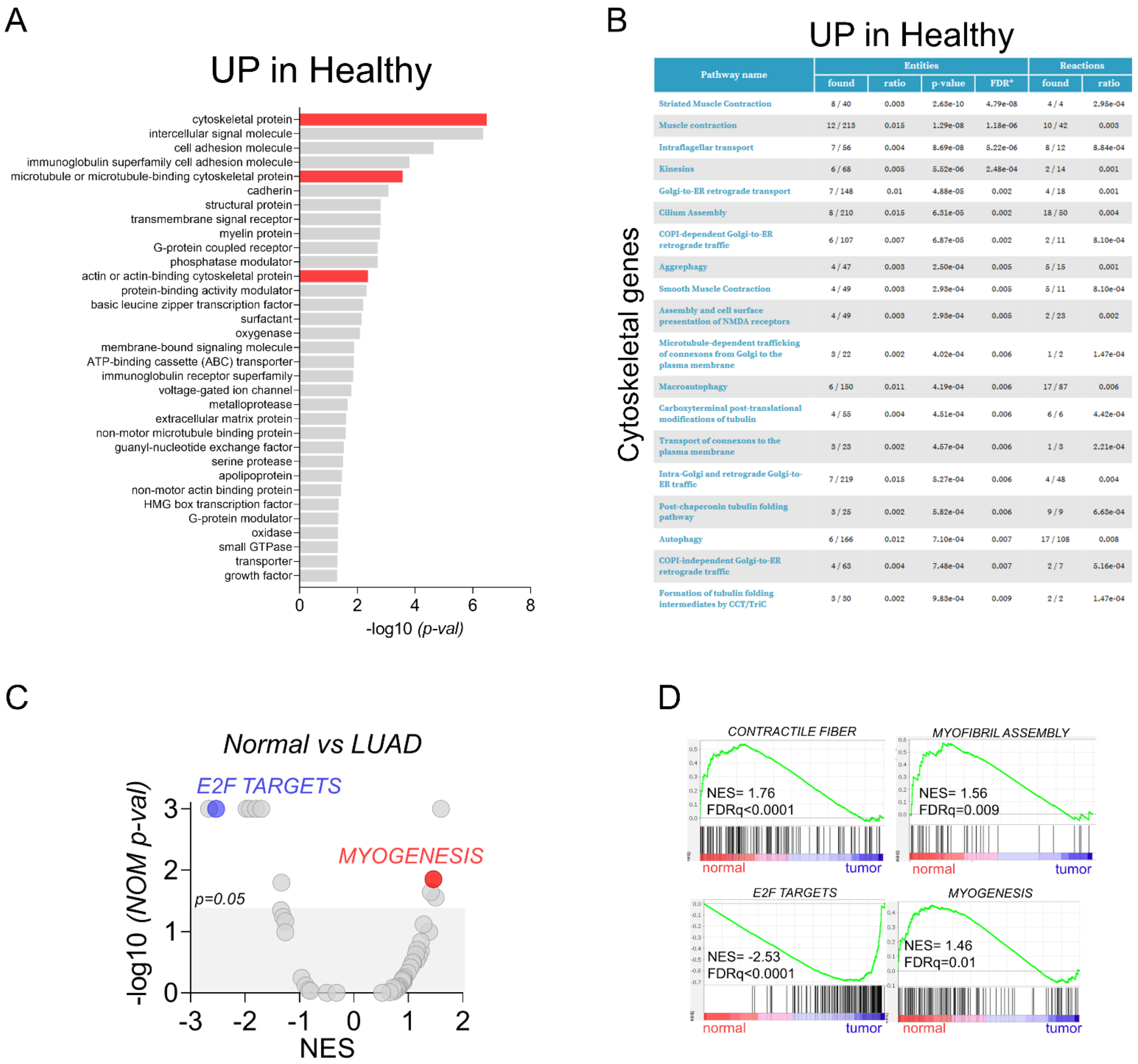


| Pathway name | Entities | | | | Reactions | |
|---|---|---|---|---|---|---|
| | found | ratio | p-value | FDR* | found | ratio |
| Striated Muscle Contraction | 8 / 40 | 0.003 | 2.63e-10 | 4.79e-08 | 4 / 4 | 2.95e-04 |
| Muscle contraction | 12 / 213 | 0.015 | 1.29e-08 | 1.18e-06 | 10 / 42 | 0.003 |
| Intraflagellar transport | 7 / 56 | 0.004 | 8.69e-08 | 5.22e-06 | 8 / 12 | 8.84e-04 |
| Kinesins | 6 / 68 | 0.005 | 5.52e-06 | 2.48e-04 | 2 / 14 | 0.001 |
| Golgi-to-ER retrograde transport | 7 / 148 | 0.01 | 4.88e-05 | 0.002 | 4 / 18 | 0.001 |
| Cilium Assembly | 8 / 210 | 0.015 | 6.31e-05 | 0.002 | 18 / 50 | 0.004 |
| COPI-dependent Golgi-to-ER retrograde traffic | 6 / 107 | 0.007 | 6.87e-05 | 0.002 | 2 / 11 | 8.10e-04 |
| Aggrephagy | 4 / 47 | 0.003 | 2.50e-04 | 0.005 | 5 / 15 | 0.001 |
| Smooth Muscle Contraction | 4 / 49 | 0.003 | 2.93e-04 | 0.005 | 5 / 11 | 8.10e-04 |
| Assembly and cell surface presentation of NMDA receptors | 4 / 49 | 0.003 | 2.93e-04 | 0.005 | 2 / 23 | 0.002 |
| Microtubule-dependent trafficking of connexons from Golgi to the plasma membrane | 3 / 22 | 0.002 | 4.02e-04 | 0.006 | 1 / 2 | 1.47e-04 |
| Macroautophagy | 6 / 150 | 0.011 | 4.19e-04 | 0.006 | 17 / 87 | 0.006 |
| Carboxyterminal post-translational modifications of tubulin | 4 / 55 | 0.004 | 4.51e-04 | 0.006 | 6 / 6 | 4.42e-04 |
| Transport of connexons to the plasma membrane | 3 / 23 | 0.002 | 4.57e-04 | 0.006 | 1 / 3 | 2.21e-04 |
| Intra-Golgi and retrograde Golgi-to-ER traffic | 7 / 219 | 0.015 | 5.27e-04 | 0.006 | 4 / 48 | 0.004 |
| Post-chaperonin tubulin folding pathway | 3 / 25 | 0.002 | 5.82e-04 | 0.006 | 9 / 9 | 6.63e-04 |
| Autophagy | 6 / 166 | 0.012 | 7.10e-04 | 0.007 | 17 / 108 | 0.008 |
| COPI-independent Golgi-to-ER retrograde traffic | 4 / 63 | 0.004 | 7.48e-04 | 0.007 | 2 / 7 | 5.16e-04 |
| Formation of tubulin folding intermediates by CCT/TriC | 3 / 30 | 0.002 | 9.83e-04 | 0.009 | 2 / 2 | 1.47e-04 |

**Supp Figure 10:**
A: Top significantly overrepresentated protein classes (pantherbd.org) in healthy lungs versus lung adenocarcinoma (log2FC>1, p-value<0.05) using TCGA database.
B: Reactome pathways associated with cytoskeletal-related genes overexpressed in healthy lungs versus lung adenocarcinoma
C: Volcano plot of the differentially regulated gene signatures between healthy lungs versus lung adenocarcinoma revealed by GSEA analysis (Hallmarks and C5).
D: GSEA analysis of contractile fiber, myofibril assembly, E2F_targets and myogenesis signatures in healthy lungs versus lung adenocarcinoma.

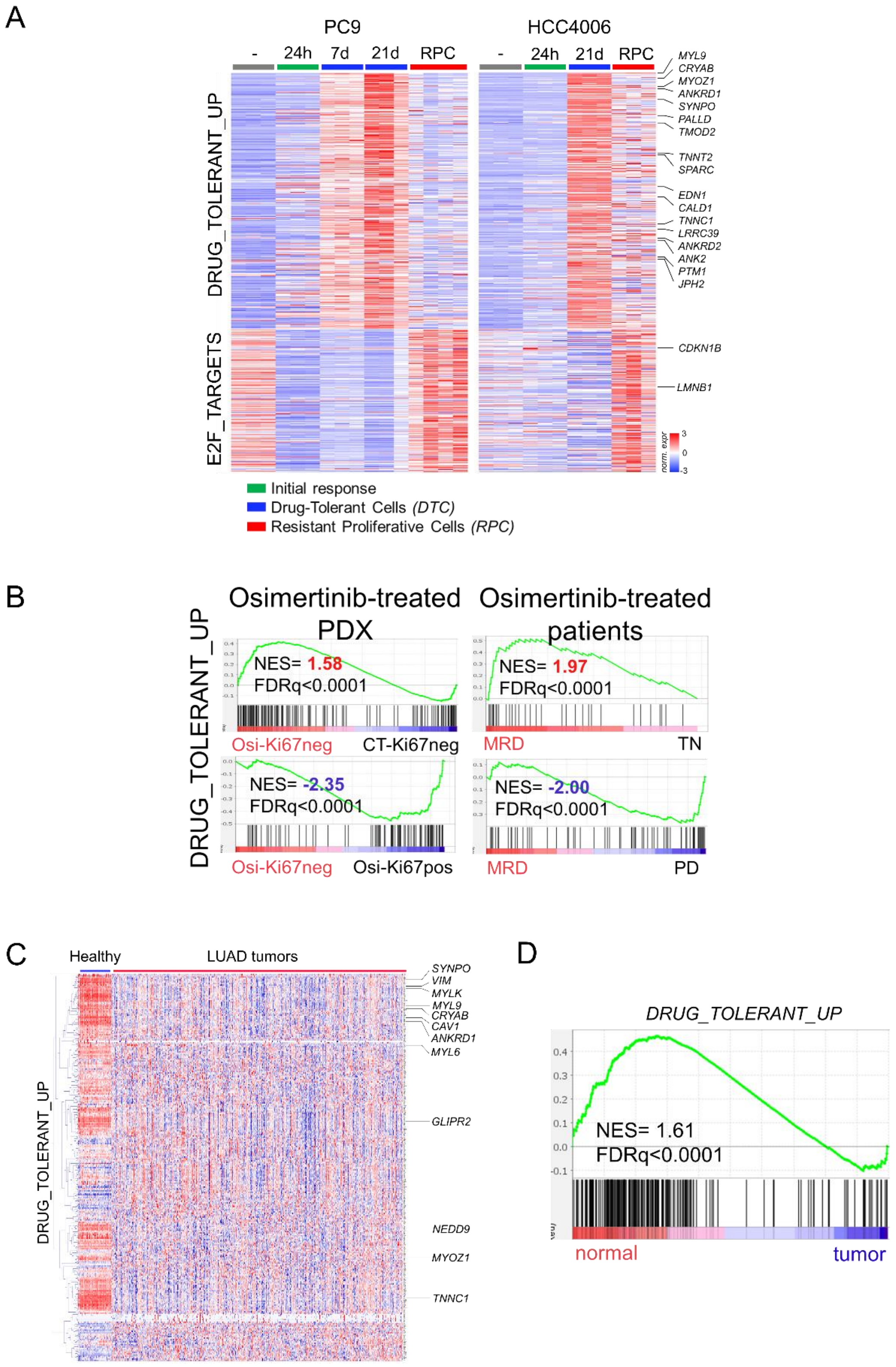


**Supp Figure 11:**

A: Heatmap of the Drug-Tolerant-associated gene signature composed of the genes that are commonly upregulated in DTC (i.e. log2FC>1, p-value<0.05 in at least 4 out of 5 cell lines).

B: Left: GSEA analysis of the drug-tolerant signature in the PDX EGFR-L858R/T790M model, comparing osimertinib-treated Ki67-negative versus vehicle-treated Ki67-negative population as a model of stable disease (top), and osimertinib-treated Ki67-positive versus osimertinib-treated Ki67-negative population as a model of relapsing tumors (bottom).

C: Heatmap of the Drug-Tolerant-associated gene signature in healthy lung and LUAD (log2FC>1, p-value<0.05) using TCGA data.

D: GSEA analysis of the drug-tolerant signature in the healthy lung versus LUAD (log2FC>1, p-value<0.05)

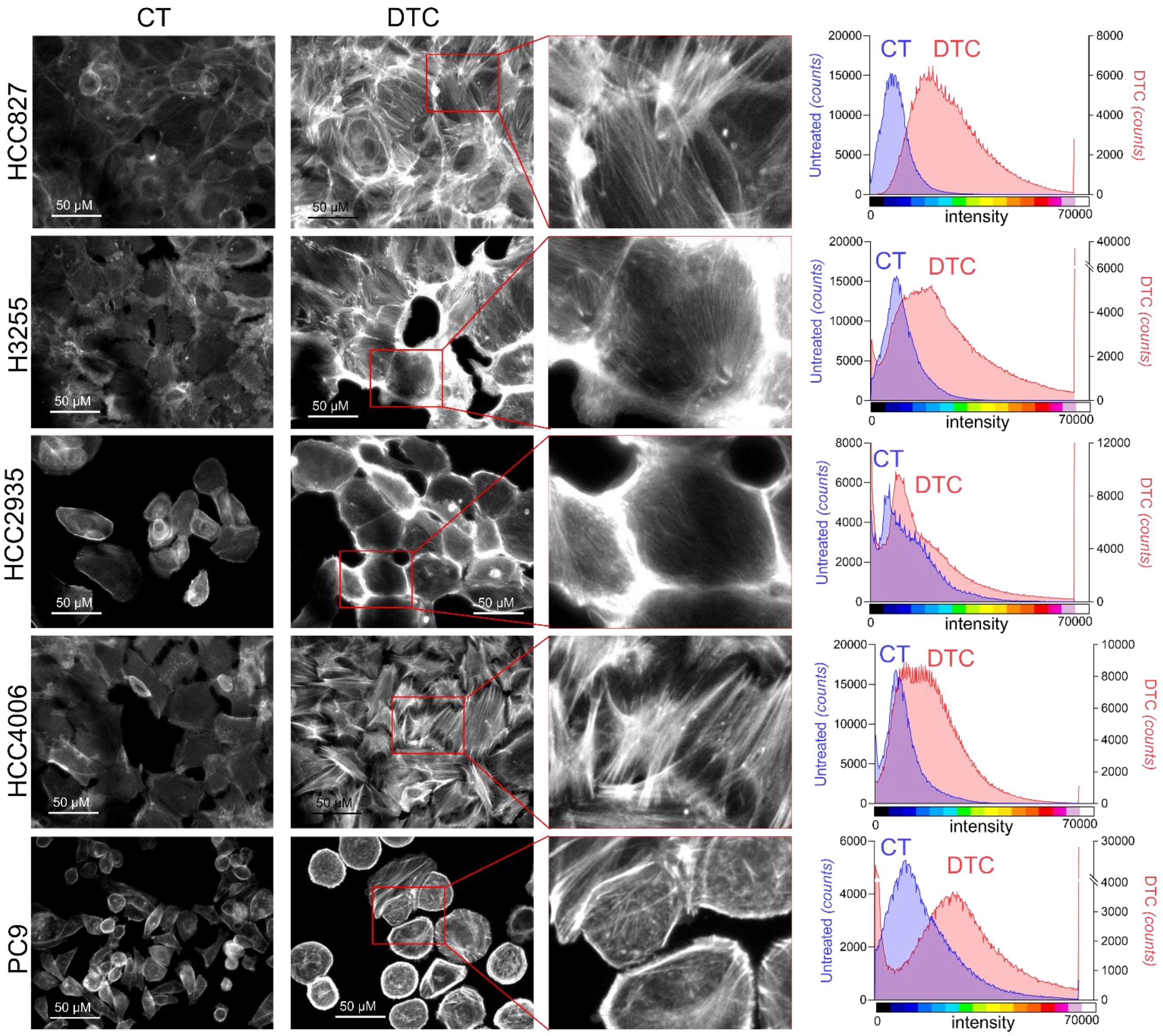


**Supp Figure 12:**
Phalloidin F-actin staining of HCC827, H3255, HCC2935, HCC4006 and PC9 control cells or treated cells with osimertinib (1µM) until DTC state (X20 magnification, scale bar = 50µM). Intensity plots were carried out by Image J software.

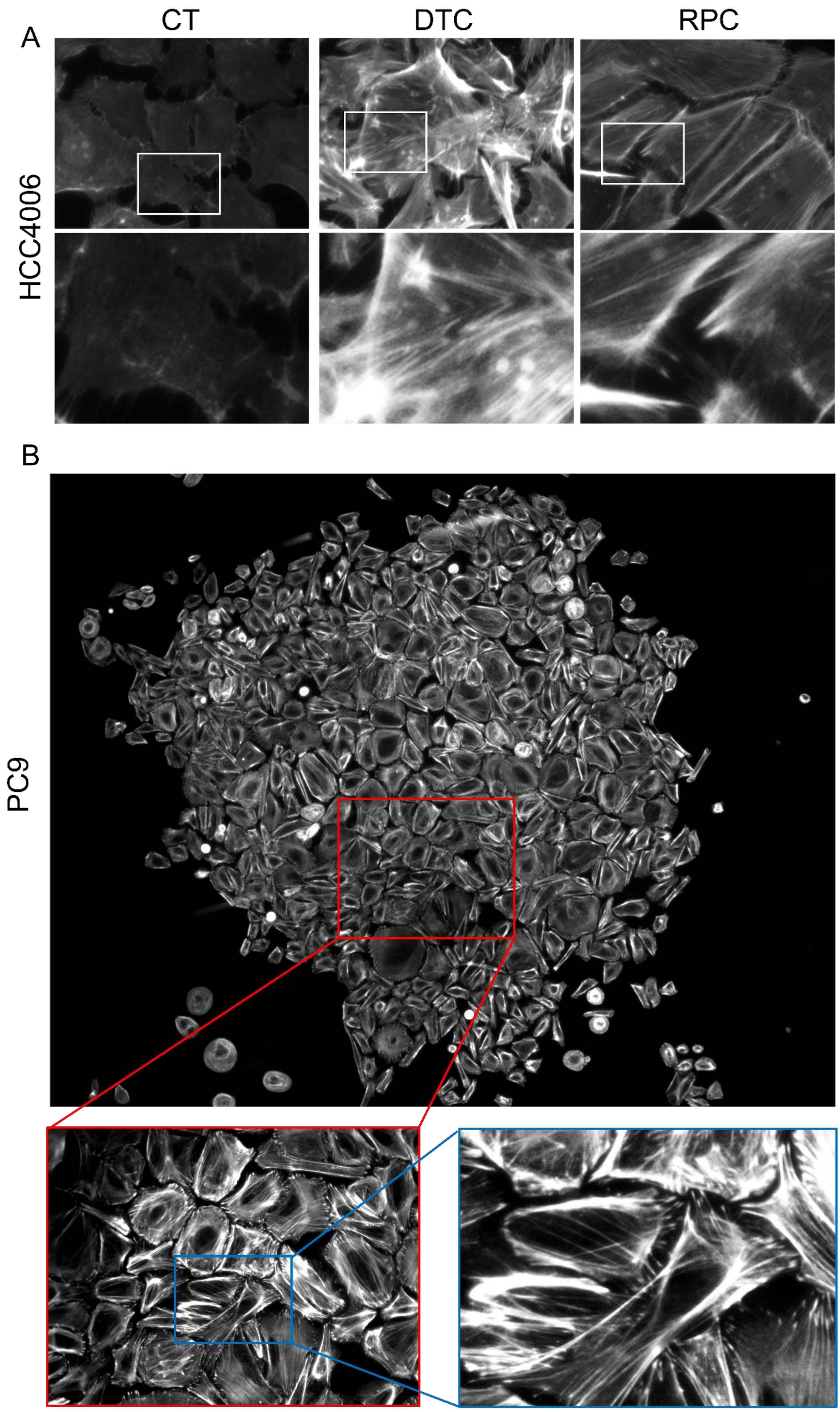


**Supp Figure 13:**
A: Phalloidin F-Actin staining of HCC4006 control cells or treated cells with osimertinib (1µM) until DTC and RPC (Resistant Proliferative Clone) state (X20 magnification).
B: Phalloidin F-Actin staining of PC9 cells treated with osimertinib (1µM) until RPC state (X20 magnification)

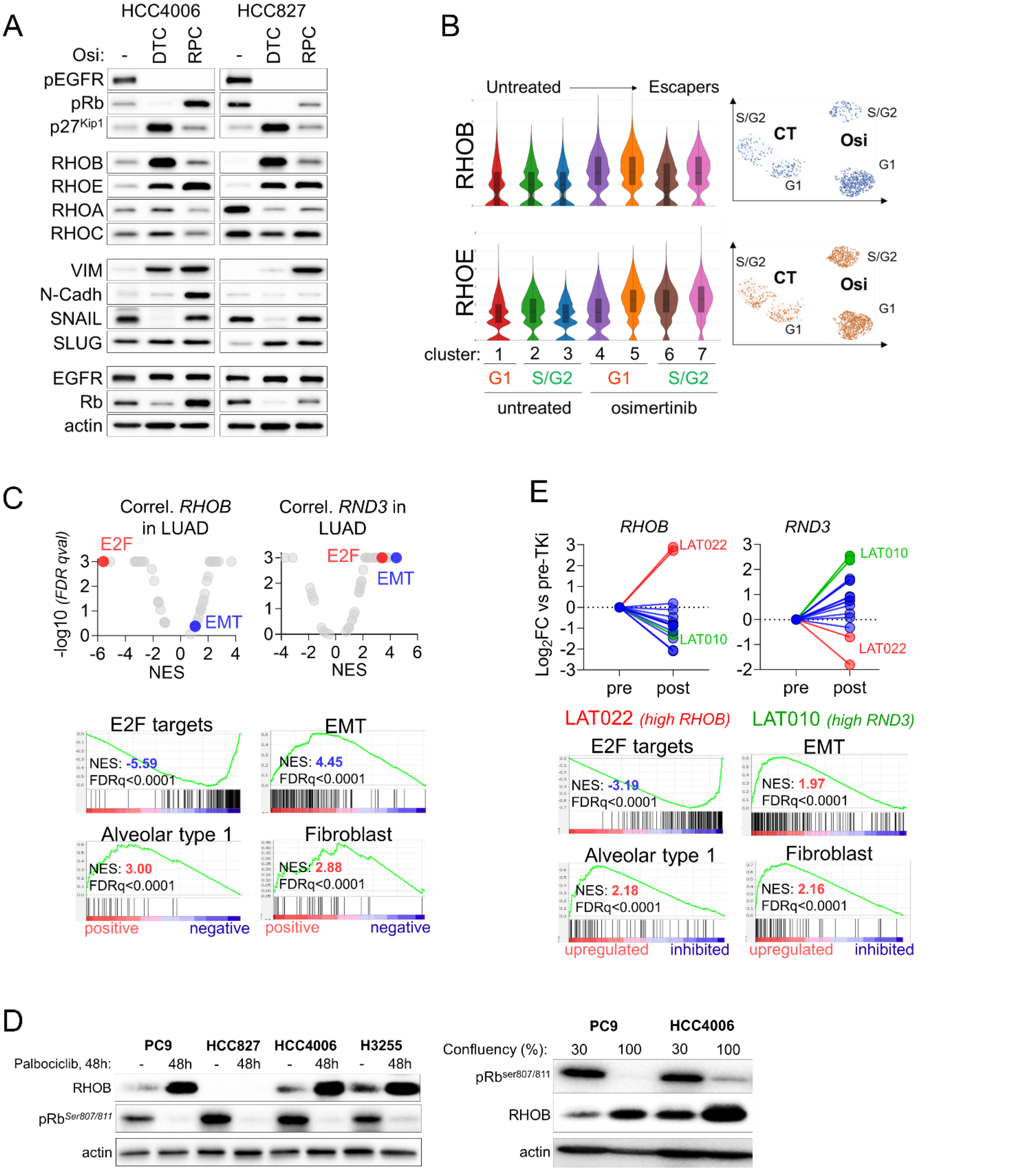


**Supp Figure 14:**

A: Protein expression by western blot of proteins related to EGFR pathway (phospho-EGFR, EGFR), cell cycle (p27Kip1, phospho-RB, RB), RhoGTPases (RHOA, RHOB, RHOC, RHOE) and EMT (Vimentin, N-cadherin, SNAIL, SLUG) on HCC4006 and HCC827 control cells or treated cells with osimertinib (1µM) until DTC or RPC state.

B: Violin plot of the Log2 mRNA expression of RHOE and RHOE (RND3) in the different clusters. UMAP representation is shown.

C: Co-expression analysis was performed using TCGA LUAD data. The list of genes that positively or negatively correlated with RHOB or RHOE mRNA expression (p-value<0.01) were analyzed by GSEA and Hallmark signatures are represented on the Volcano plot (top) and ranked GSEA plots (bottom).

D: Protein expression by western blot of phospho-RB and RHOB on PC9, HCC827, HCC4006, H3255 control cells or treated cells with palbociclib (CDK4/6 inhibitor, 1µM) for 48 hours. Protein expression by western blot of phospho-RB and RHOB on PC9, HCC4006 cells with 30% or 100% of confluency.

E: Top: Evolution of RHOB and RHOE (RND3) mRNA expression levels between pre-treatment and first progression to osimertinib (data from Roper et al, Cell Rep Med, 2020). Bottom: GSEA analysis of E2F_targets, AT1, EMT and fibroblast signatures in resistant versus pre-treated tumors. LAT022 patient displayed a low proliferation rate at relapse, suggesting that tumor sample used for RNAseq was still responding to osimertinib treatment. LAT010 patient displayed a high EMT score at relapse which correlated with higher RND3 expression.

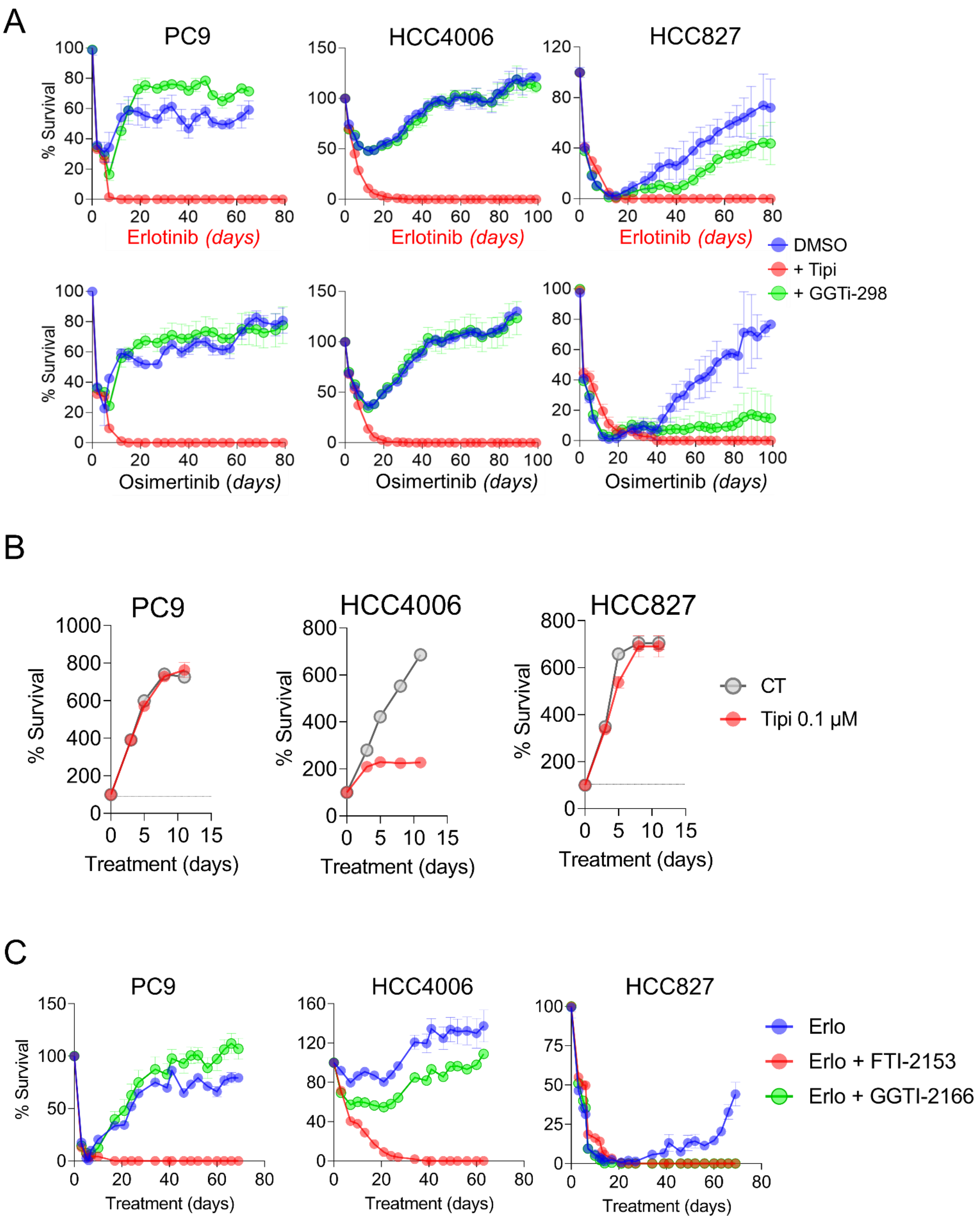


**Supp Figure 15:**

A: Cell survival (%) of PC9, HCC4006 and HCC827 cells treated with erlotinib or osimertinib (1 μM) alone or in combination with tipifarnib (1μM) or GGTi-298 (1μM).

B: Cell survival (%) of PC9, HCC4006 and HCC827 control cells or treated with tipifarnib (0.1μM).

C: Cell survival (%) of PC9, HCC4006 and HCC827 cells treated with erlotinib (1 μM) alone or in combination with FTI-2153 (1μM) or GGTi-2166 (1μM).

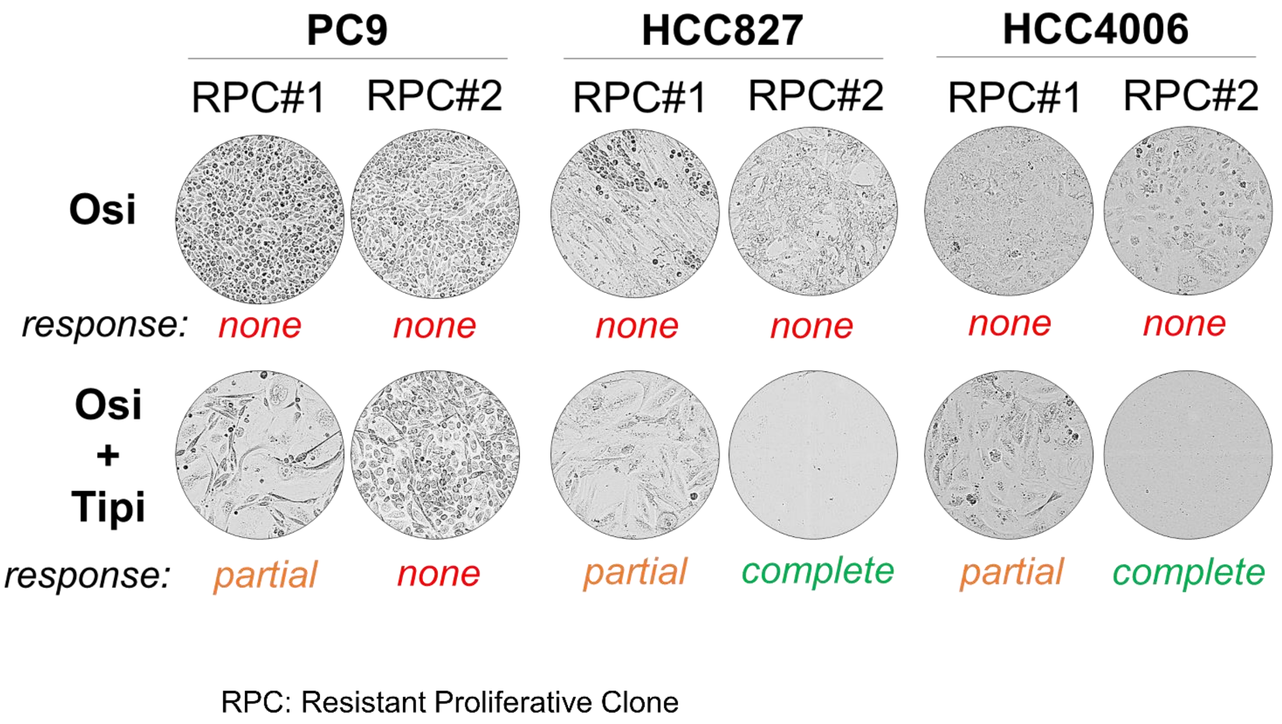


**Supp Figure 16:**
Phase contrast images of PC9, HCC827, HCC4006 osimertinib-RPC cells treated with osimertinib 1µM alone or in combination with tipifarnib 1µM.

Figarol *et al.*

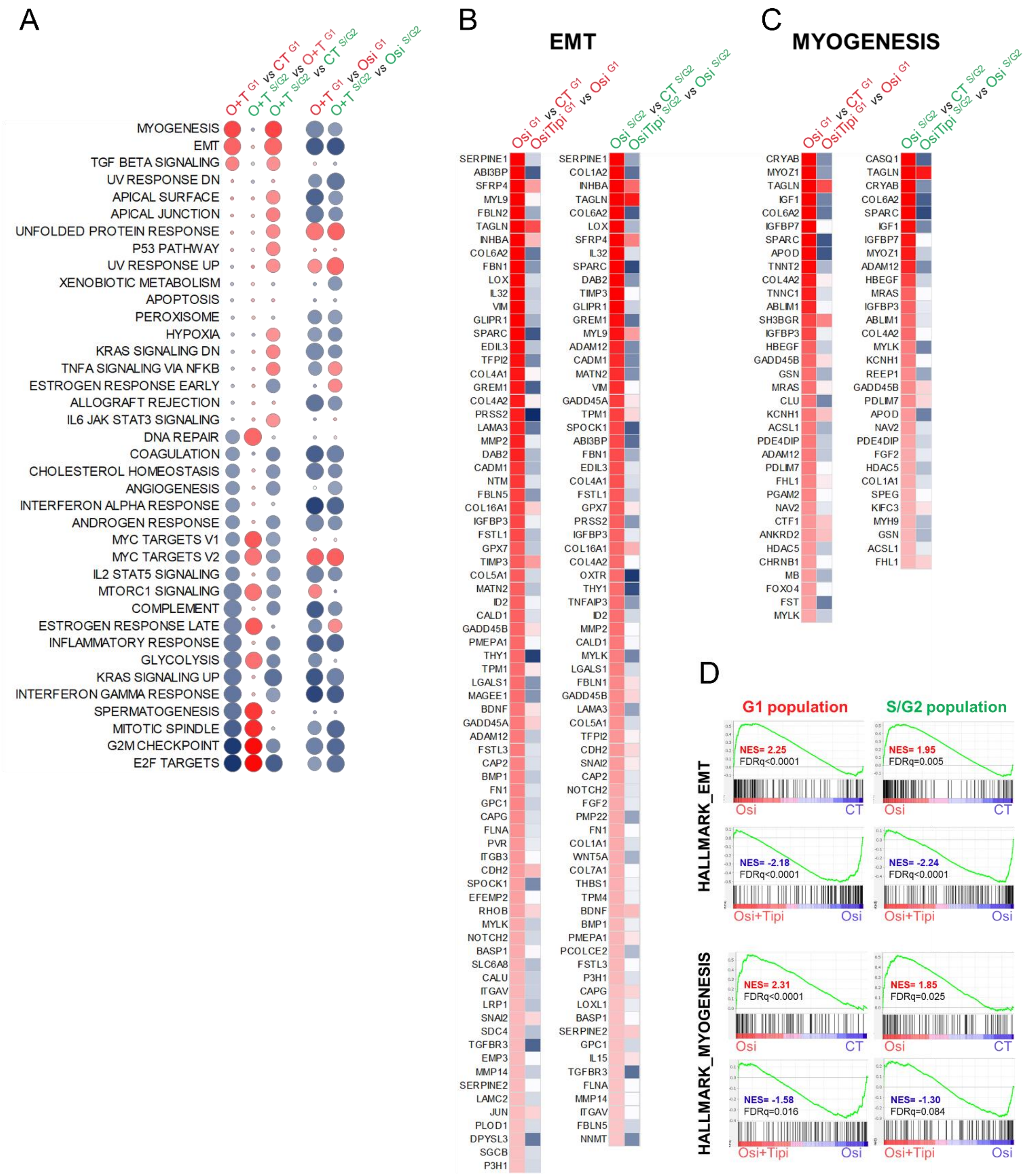


**Supp Figure 17:**
A: GSEA of Hallmark gene sets in HCC4006 for indicated cell populations.
B, C. Differential mRNA expression of EMT-related genes (B) and myogenesis-associated gene (C) for indicated cell populations.
D: GSEA analysis of EMT and myogenesis signatures in osimertinib (Osi) or osimertinib+Tipifarnib-treated HCC4006 cells.

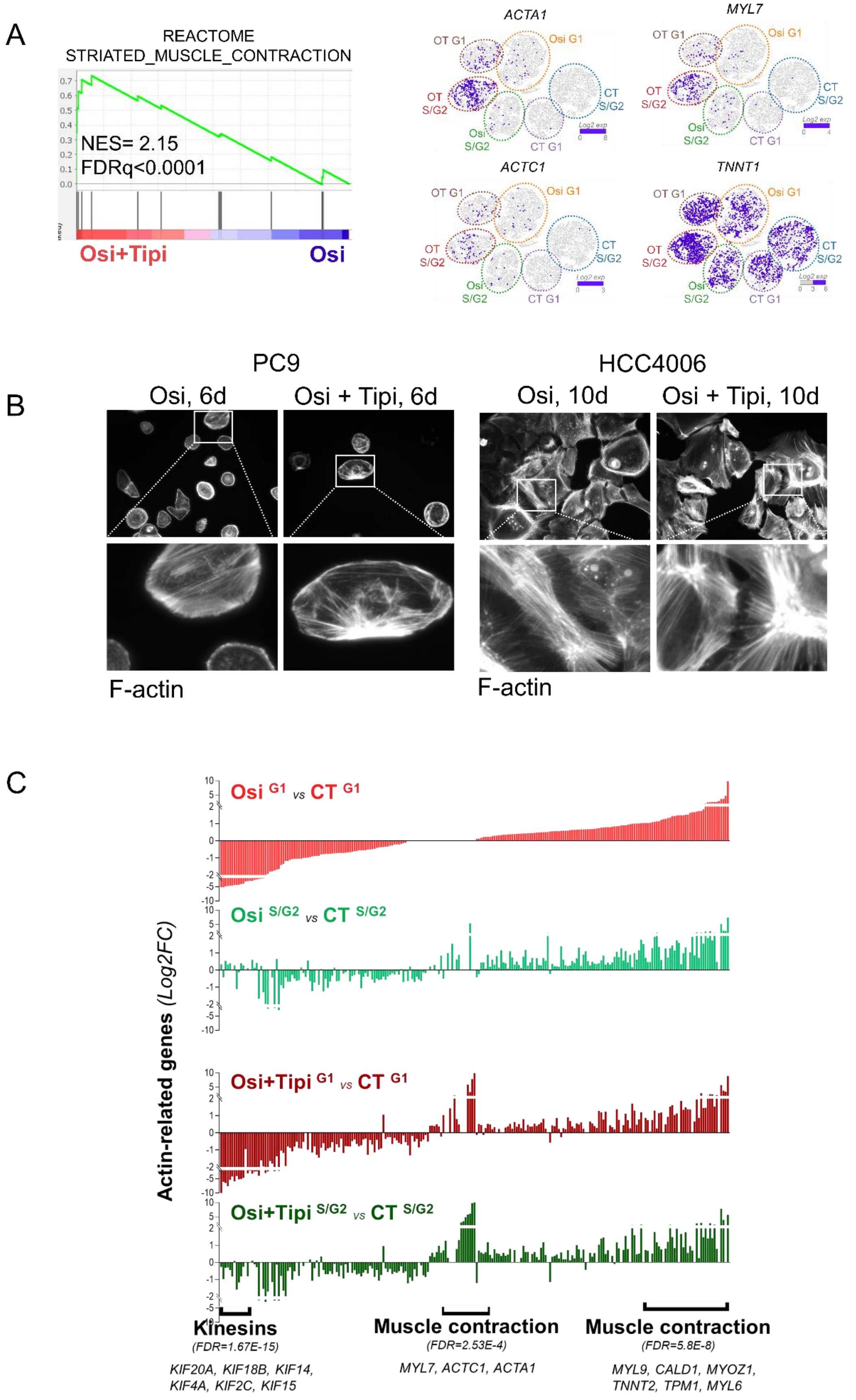


**Supp Figure 18:**

A: GSEA analysis of striated muscle contraction signatures in HCC4006 clonal cells treated with osi+tipi versus osi (left). Right: t-SNE representation of muscle-associated genes ACTA1, MYL7, ACTC1 and TNNT1.

B: Phalloidin F-Actin staining of PC9 and HCC4006 cells treated with osimertinib (1µM) alone or in combination with tipifarnib (1µM) for 10 days (X20 magnification).

C: Log2 fold change of actin-related genes expression in indicated-treated HCC4006 clonal cells.

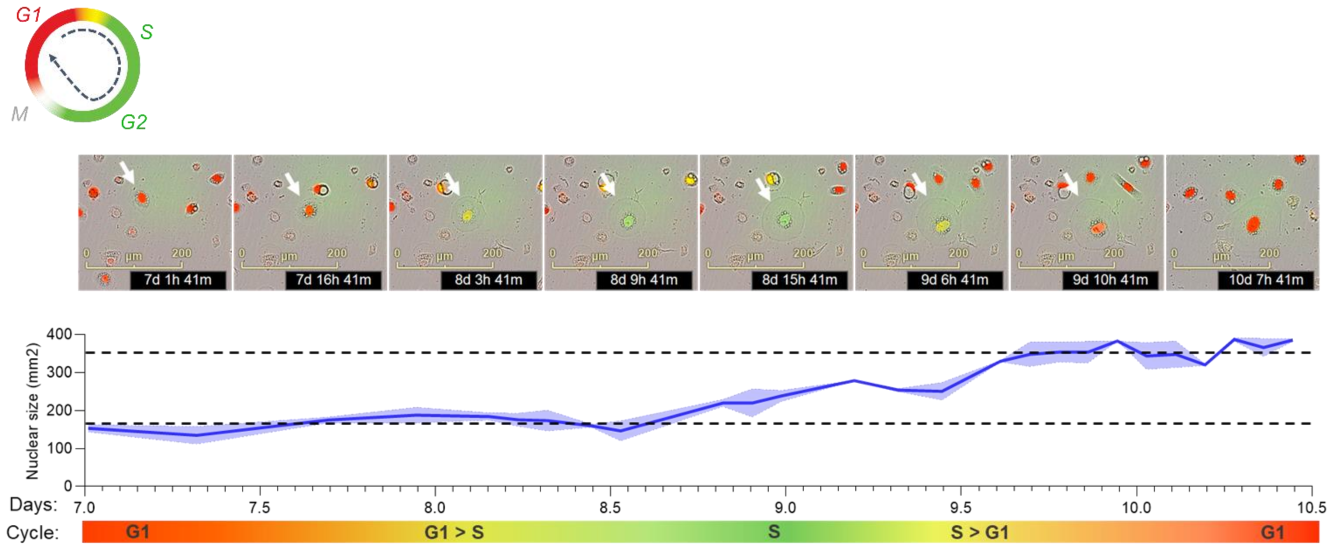


**Supp Figure 19:**
Top panel: Images of a PC9 subclonal cell under osimertinib (1 µM) treatment showing endoreplication process. Bottom panel: Quantification of nuclear size of the same PC9 subclonal cell under osimertinib (1 µM) treatment.

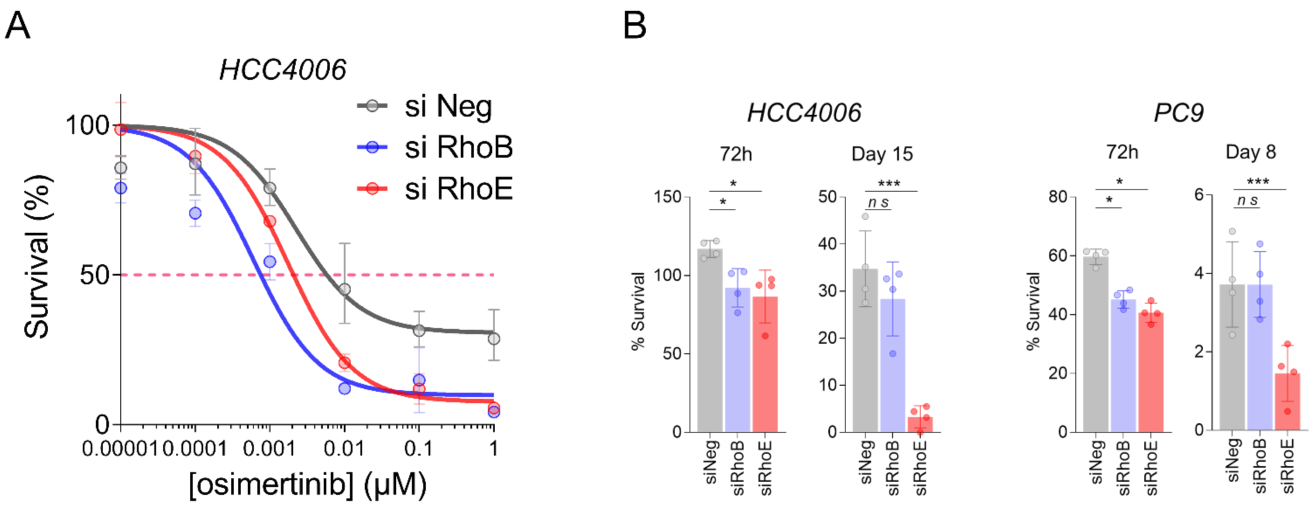


**Supp Figure 20:**
Cell survival by cytotoxicity assay of HCC4006 cells transfected with siRNA control (Neg) or targeting RHOB or RHOE under osimertinib treatment for 5 days.

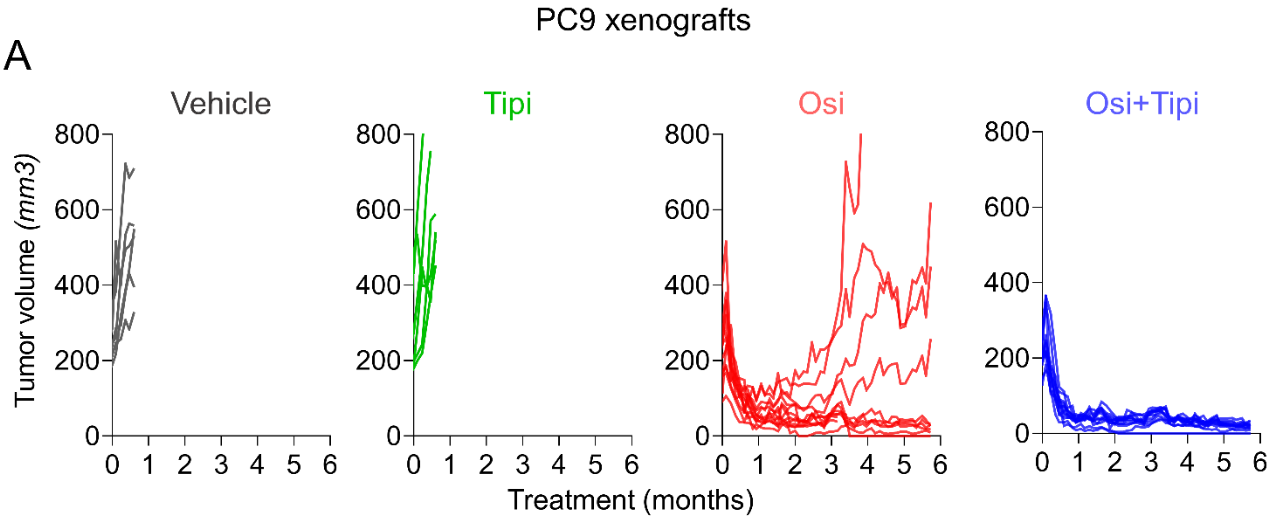


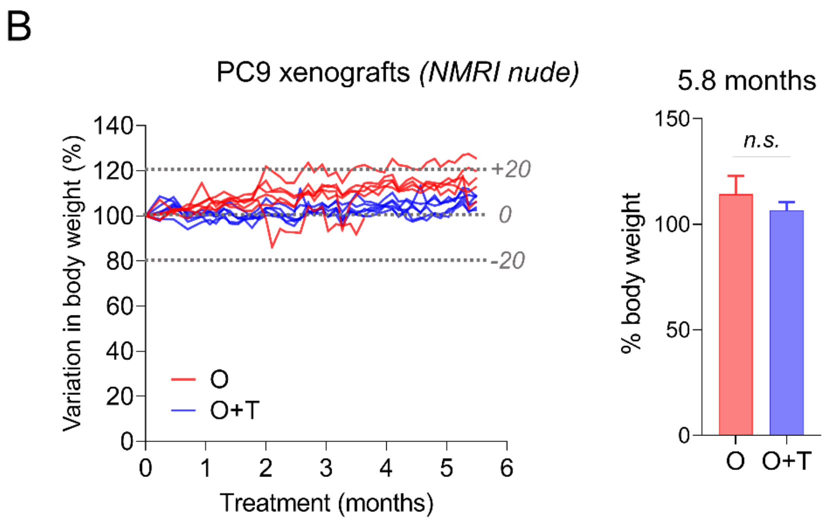


**Supp Figure 21:** PC9-xenografts studies.
A. Individual tumor volume (mm3) treated 5 days/week with vehicle, Tipifarnib (Tipi, 80mg/kg, b.i.d.), Osimertinib (Osi, 5 mg/kg, q.d), or by the combo (Osi + Tipi). n = 6 tumors in the vehicle and Tipifarnib arms, and n = 10 in the Osimertinib arm and n=12 in the combination arms.
B. Variation of body weight (%) along treatment course (left) and at 5.8 months (right) for 5 mice treated with Osimertinib (5 mg/kg) and 6 mice treated with the combination of Osimertinib and Tipifarnib (80 mg/kg).

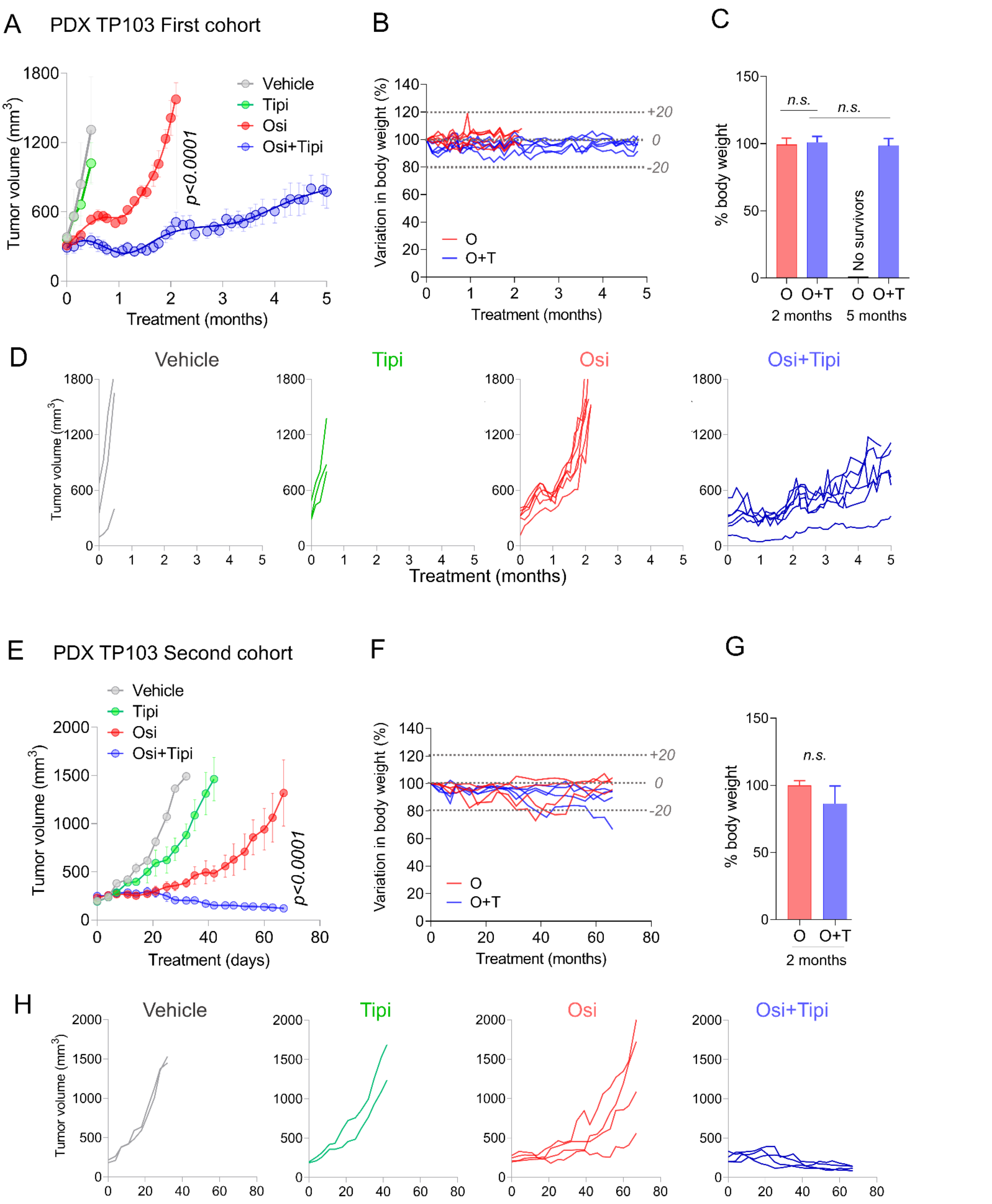


**Supp Figure 22:**
Analysis of treatment efficacy in an EGFR-L858/T790M lung adenocarcinoma PDX model (TP103). A-D. First cohort of mice. E-H. Second cohort of mice. Mean ± SEM (A, E) and individual (D, H) tumor volume (mm3) treated 5 days/week with vehicle, Tipifarnib (Tipi, 80mg/kg, b.i.d.), Osimertinib (Osi, 5 mg/kg, q.d), or with the combo (Osi + Tipi). Variation in body weight (%) during treatment (B, F) and at the end of treatment (C, G) with Osimertinib and the combination treatments. For the first cohort (A-D) n = 3 tumors in the vehicle and Tipifarnib arms and n = 6 tumors in the Osimertinib and combination arms. For the second cohort (E-H) n = 2 tumors in the vehicle and Tipifarnib arms and n = 4 tumors in the Osimertinib and combination arms.

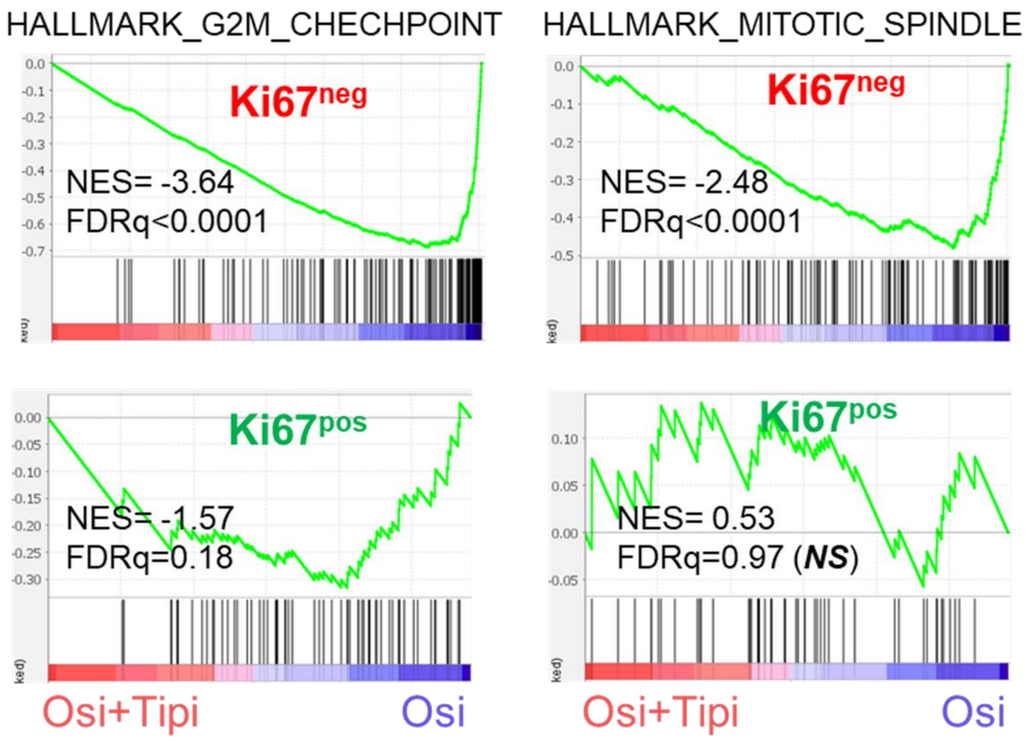


**Supp Figure 23:**
GSEA analysis of H_G2M_checkpoint and mitotic_spindle signatures in Osimetinib+Tipifarnib versus Osimertinib-treated EGFR-mutated PDX
